# Rewiring of RSK-PDZ interactions by linear motif phosphorylation

**DOI:** 10.1101/419721

**Authors:** Gergo Gogl, Beata Biri-Kovacs, Fabien Durbesson, Pau Jane, Yves Nomine, Camille Kostmann, Viktoria Bilics, Marton Simon, Attila Remenyi, Renaud Vincentelli, Gilles Trave, Laszlo Nyitray

## Abstract

Protein phosphorylation is a key regulator of protein-protein interactions. How does the interactome of a protein change during extracellular stimulations? While many individual examples of phosphorylation-regulated interactions were described previously, studies addressing the interactome changes induced by a particular phosphorylation event remain scarce. Here, we try to answer this question, by focusing on interactions between a phosphorylable PDZ-binding linear motif and the entire complement of human PDZ domains. Using a combination of *in vitro* quantitative techniques and cell-based approaches, we demonstrate that the activation of the mitotic effector kinase RSK1 causes dramatic changes in its connectivity with PDZ domain containing proteins. These changes consist of modulations of the binding affinity of numerous interactions, rather than on/off switching of a few interactions. Our results highlight the previously unappreciated role of phosphorylation in the complex and subtle rewiring of large numbers of protein-protein interactions.

## BACKGROUND

Within a given organism, the size of the protein-protein interactome significantly exceeds the size of the proteome (Gould *et al*, 2010 Moreover, the interactome is not static. A key driving force behind its dynamic nature is the fine tuning of individual interactions by reversibly altering their biochemical properties (e.g. binding affinity). A large proportion of protein-protein interactions (PPIs) is mediated by families of globular domains binding to intrinsically disordered short interaction segments, often referred to as docking peptides or short linear motifs (Neduva & Russell, 2006). While these linear motif-based interactions can be regulated in many ways, the most universal mechanism appears to be reversible protein phosphorylation (Neduva & Russell, 2005).

An interesting instance of phosphorylation-based regulation of domain-motif interactions are those involving PDZ domains. PDZ domains belong to one of the most common family of globular domains, with 266 members in the human proteome (Luck *et al*, 2012). They recognize short linear motifs called PBMs (for “PDZ-Binding Motifs”) at the extreme C-terminus of their target proteins. PBMs systematically contain a hydrophobic residue (most frequently Val or Leu) at their C-terminus (numbered as position 0) and are classified in three main classes based on the residue at position -2 (Ser/Thr in the most common class 1, hydrophobic in class 2 and acidic in class 3) (Sheng & Sala, 2001).

PDZ-PBM interactions are involved in various cellular processes, and are especially common in intracellular signaling pathways. For example, all isoforms of the ribosomal S6 kinase (RSK) of the MAPK pathway contain a functional class 1 PBM (Thomas *et al*, 2005). RSK has an emerging role in multiple cancer types such as glioblastoma or melanoma (Sulzmaier *et al*, 2016) (Hartman *et al*, 2014). Upon mitogenic stimuli, a series of phosphorylation events leads to the activation of the MAP kinase ERK1/2 (Pouysségur *et al*, 2002). RSK is one of the strongest interaction partner of ERK and its complex activation mechanism is also initiated by ERK phosphorylation (Figure 1A) (Alexa *et al*, 2015). The C-terminal tail of RSK is a multifunctional linear motif as it contains (partially overlapping) binding sites for ERK, S100B, a tyrosine kinase, phosphatase(s) and PDZ domains (Garai *et al*, 2012) (Gógl *et al*, 2015) (Kang *et al*, 2008). Additionally, activated RSK will autophosphorylate its own PBM within its intrinsically disordered tail, which will probably affect all of these interactions (Gógl *et al*, 2018). The RSK1 PBM contains three potential autophosphorylation sites, while other isoforms contain only two (Figure 1B), and among these, the major site can be found at the -3 position (Dalby *et al*, 1998). The consequence of the major autophosphorylation on PDZ binding is not known, however we have some clues. For example, Thomas et al. showed that no change could be observed with RSK1/2 phosphomimics (at -3) in the interaction with MAGI1, SHANK1 or GRIP1, and they suggested that both inactive and active RSKs likely bind to PDZ domain proteins (Thomas *et al*, 2005). Similarly, our recent work showed that phosphorylation of RSK1 only mildly changed the interaction with MAGI1 (Gógl *et al*, 2018). In contrast, a recent publication revealed that the phosphorylation (or phosphomimics) at the analogous site triggered the association between RSK1/3 and the PDZ domain of SCRIBBLE and abolished the interaction between RSK3 and the PDZ domain of SHANK1 (Sundell *et al*, 2018). These results indicated that RSK activation might induce a complex reshuffling of its PDZ domain mediated interactome.

**Figure 1.**
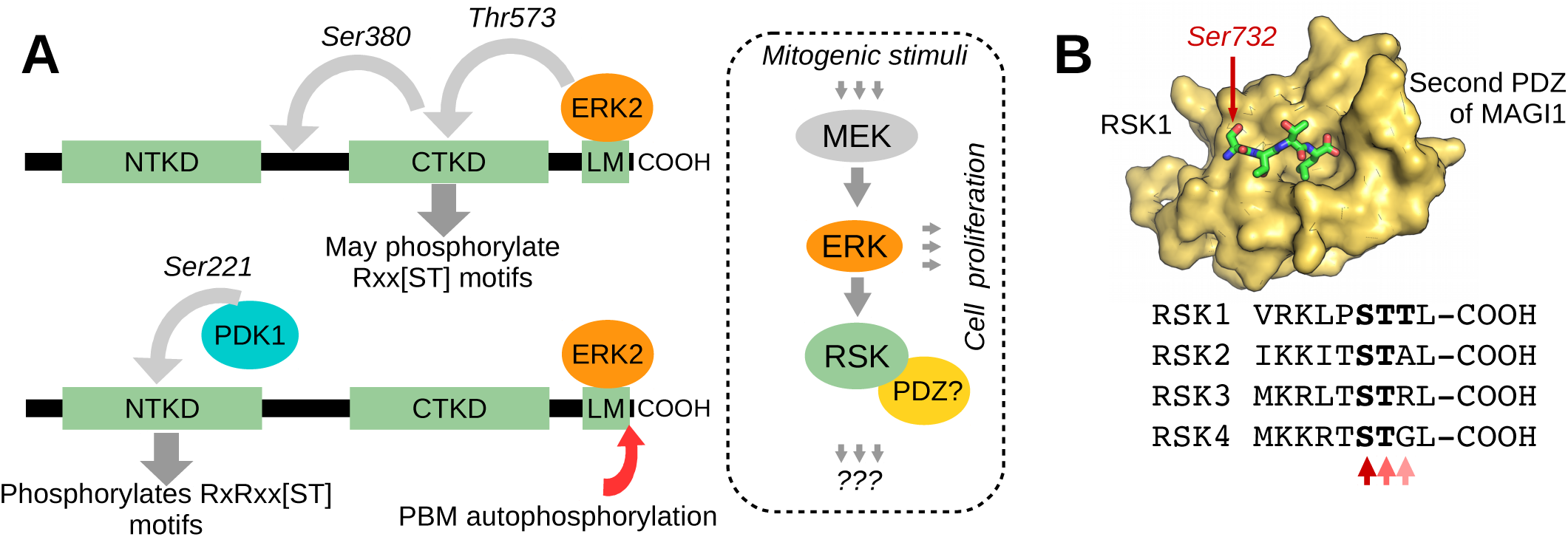
The activation of RSK includes a feedback phosphorylation site, which can affect PDZ binding. (A) Activation of the tandem kinase RSK is a multi-step process. Activation of RSK is initiated by ERK docking, which is followed by the phosphorylation of the C-terminal kinase domain (CTKD) (Alexa *et al*, 2015). The active CTKD will phosphorylate a linker site between the kinase domains, which will create a docking motif for PDK1 (Frödin *et al*, 2002). In the end, PDK1 will activate the N-terminal kinase domain (NTKD) (Frödin *et al*, 2000). Usually, only the NTKD is considered as an effector kinase and the CTKD is only associated with a self-regulatory role and one of these activated kinases will phosphorylate its C-terminal PBM. While RSK is an effector of the mitogenic ERK pathway, we have limited information about its intracellular role. (B) Each RSK isoform contains a functional, class 1 PBM. RSK1 contains 3, mutually exclusive autophosphorylation sites (at the -1,-2,-3 positions) and the other isoforms contain only two (at the -2,-3 positions), but only the -3 site (Ser732 in RSK1) is considered as a major feedback site (Hornbeck *et al*, 2015). The structural panel shows the RSK1 binding to the second PDZ domain of MAGI1.

In principle, the general consensus sequence determining a PBM allows for the presence of potentially phosphorylable residues at any position except the hydrophobic C-terminal position (Saras & Heldin, 1996). Therefore PDZ-PBM interactions are an ideal models to study the impact of phosphorylation on PPIs. A first challenge in such a study would be to analyze the effect of phosphorylation of a given PBM on its binding to all its putative partners in the human proteome, i.e. on all known human PDZ domains. A second challenge would be to use a quantitative method based on binding affinity measurements, rather than a simplistic binary description of the binding mode (“bind” or “do not bind”). Indeed, a recently developed “holdup assay” allows one to measure the individual binding affinities of the 266 known human PDZ domains for a single PBM, thereby generating a “quantitative PDZome-binding profile” of the motif (Vincentelli *et al*, 2015). Here, we applied this holdup assay method to generate the PDZome-binding profiles of both the unphosphorylated and phosphorylated RSK1 PBMs. Next, the strongest PDZ-PBM interactions were confirmed and further characterized by using low-throughput biophysical approaches. Several interactions were also confirmed with full-length proteins in mammalian cells, by using an intracellular luciferase fragment complementation assay. Moreover, in this assay, EGF stimulation of the cells, known to induce autophosphorylation of RSK1 PBM, modulated the binding of full-length RSK1 to its target proteins in a way that was consistent with our *in vitro* observations.

## RESULTS

### PDZome-binding profiles of native and phosphorylated RSK1 PBMs

To investigate how phosphorylation can modulate the binding of the PBM of RSK to PDZ domains, we used the automated high-throughput holdup assay, which allows to measure binding intensities (BI values) for large numbers of domain-motif pairs experimentally. As compared to the original work describing this approach (Vincentelli *et al*, 2015), we used an updated version of our PDZ library, which now includes the 266 known human PDZ domains (Duhoo *et al*). Here, out of 266 PDZ, we could quantify the interaction of 255 PDZ for the unphosphorylated RSK1 peptide and 252 for the phosphorylated form. Both data sets were plotted in the form of “PDZome-binding profiles” (Figure 2A) representing all the individual “binding intensities” (BIs) measured for each PDZ domain for the unphosphorylated and phosphorylated RSK peptides, respectively. Using BI = 0.2 as the minimal threshold for a significant PDZ-peptide interaction, the holdup assay identified 34 potential RSK1 binders, including 26 PDZ binders for the unphosphorylated peptide and 25 binders for the phosphopeptide (Figure 2, S1, Table S1). The general distribution of the PDZome-binding profiles were similar in both cases. However, phosphorylation decreased the maximal and average BIs from 0.77 to 0.54, and from 0.42 to 0.33, respectively. Furthermore, the order of the PDZ domains that bind best to the unphosphorylated and phosphorylated RSK1 PBM was markedly different, as visually illustrated by the global reshuffling of their respective profiles (Figure 2A). Taking BI > 0.2 as a threshold for significant binding, the phosphorylated RSK1 PBM lost 12 of the binding partners and gained 10 new potential binding partners as compared to the unphosphorylated peptide. This implies that approximately 35% of the potential binders interact (often with variable affinities) to both RSK1 peptides while 65% of them bind detectably to only one or the other forms of the RSK1 PBM.

**Figure 2.**
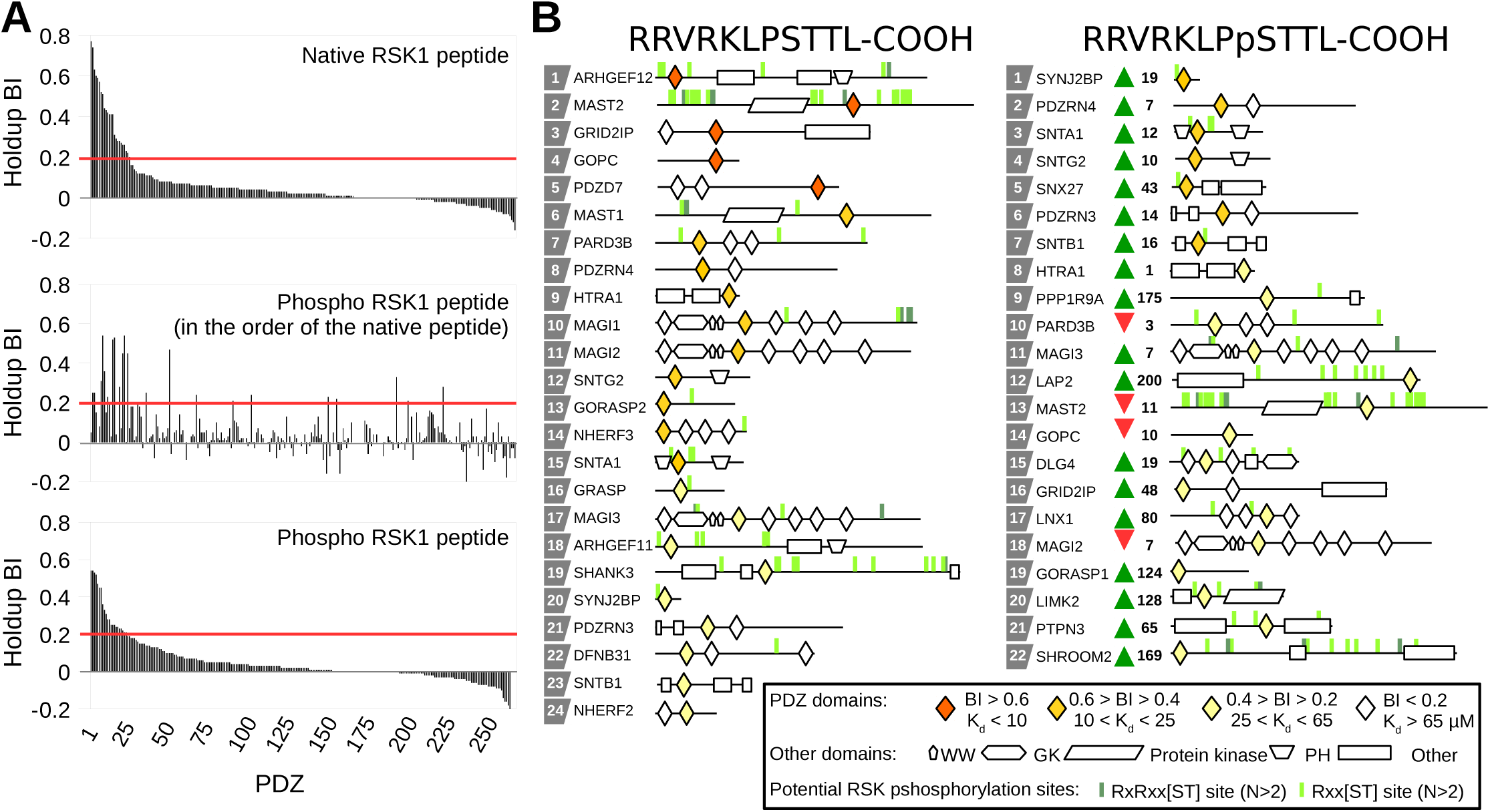
Results of the holdup assay. (A) PDZome binding profiles of unphosphorylated and phosphorylated RSK1 PBMs. All profiles use biotin as a control. A red line indicates the cutoff for a significant PDZ-PBM interaction (BI>0.2). PDZ domains in the upper and lower plots are ranked on the basis of their BIs for the indicated peptide. In the middle plot, PDZ domains are ranked on the basis of their BIs for the unphosphorylated peptide, while the plotted BI values are those obtained for the phosphopeptide. Note the considerable reshuffling of binding targets induced by phosphorylation. (B) Domain architecture of the identified interaction partners. The PDZ domains are colored according to the measured BI values. Potential RSK phosphorylation sites are highlighted in the schematic maps with sticks. Dark and light green sticks represent ideal and non ideal RSK phosphorylation sites, respectively. Phosphorylation sites were extracted from the phosphosite (Hornbeck *et al*, 2015) (an N > 2 filter was applied on the database).

The assay identified ARHGEF12 as the strongest interaction partner of the unphosphorylated peptide (Figure 3 and Table 1, Table S1). This protein is a RhoA GEF and it has recently been reported that its interaction with RSK2 is essential in RhoA activation and this interaction leads to increased cell motility in the U87MG glioblastoma cell line (Shi *et al*, 2018). We also identified strong interaction with MAST2, which is an AGC kinase similarly to RSK (Pearce *et al*, 2010). The previously characterized interaction between MAGI1 and RSK1 was found among the top binders of the unphosphorylated dataset and interestingly, our approach shows that the phosphorylation strongly disrupts the interaction in contrast to earlier works (Thomas *et al*, 2005) (Gógl *et al*, 2018). The strongest interaction partners of the phosphorylated PBM were three signal transducing adaptor proteins SYNJ2BP, SNTA1 and the E3 ubiquitin ligase PDZRN4 (Figure 3 and Table 1, Table S1).

**Figure 3.**
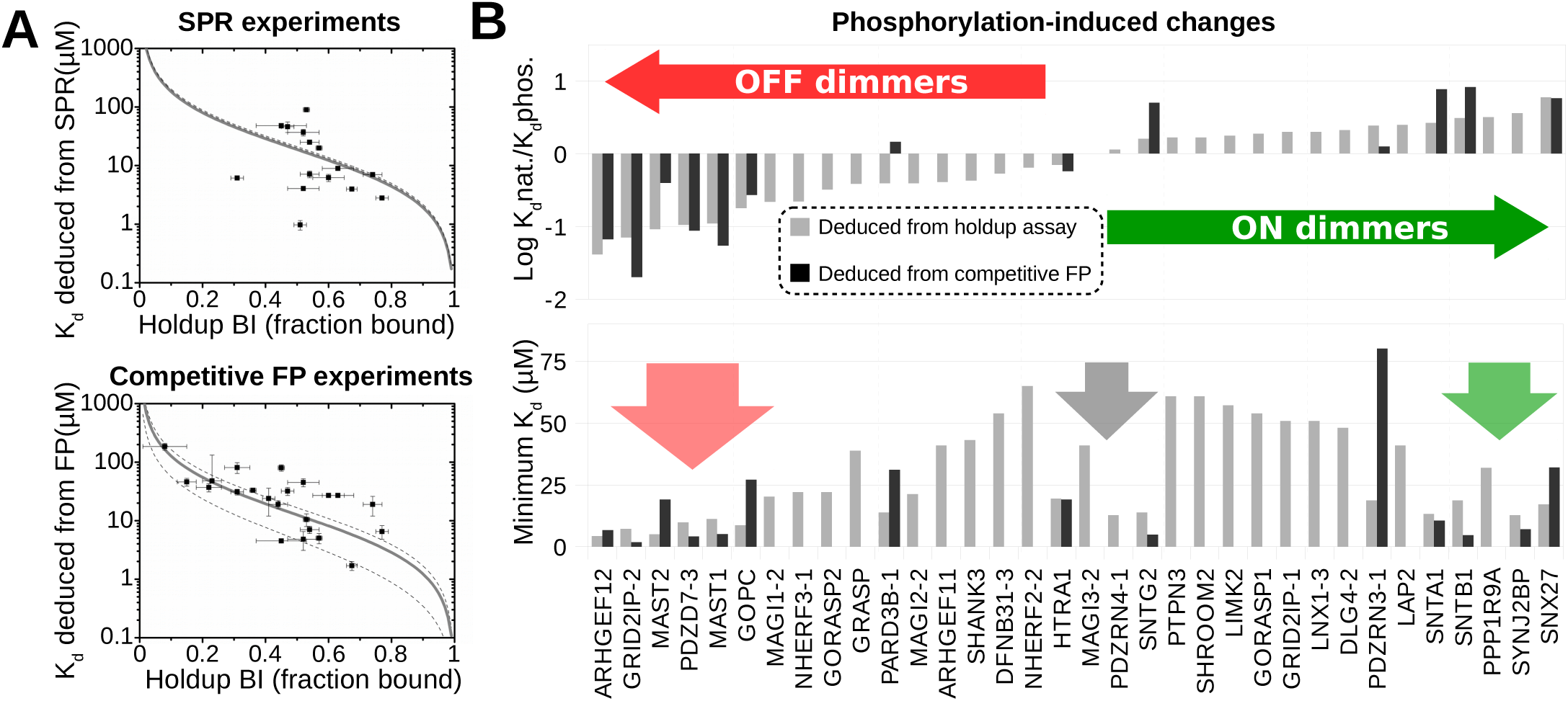
*In vitro* validation of the holdup assay. (A) Comparing orthogonal binding data obtained by holdup assay, SPR and competitive fluorescence polarization (FP). The correlation of binding intensities (BI) obtained by holdup assays to affinity constants deduced from SPR or competitive FP was fitted using a Monte Carlo approach. Despite independent fitting procedures, a similar correlation was observed in both cases. The fitting procedure delivers a value for the peptide concentration in the holdup assay. By combining the peptide concentration obtained in that way, with the concentrations of free and peptide bound PDZ domain (both delivered by the holdup assay); one can then estimate from the holdup data the dissociation constant of all human PDZ domains that interacted detectably with the RSK1 peptides. (B) Phosphorylation induces a complex rearrangement in the RSK1 PDZ interactome. Instead of two definite classes (ON or OFF switching), a continuum (ON or OFF dimming) was measured in the phospho-induced K_d_ differences. Dark gray columns show the experimentally determined K_d_ differences from the competitive FP measurements. The lower panel shows the strongest affinity (minimal K_d_) observed with either the native or phosphorylated RSK1 peptide in the same order as in the upper panel. Significant interaction partners can be found in all regions of the observed continuum.

**Table 1.**
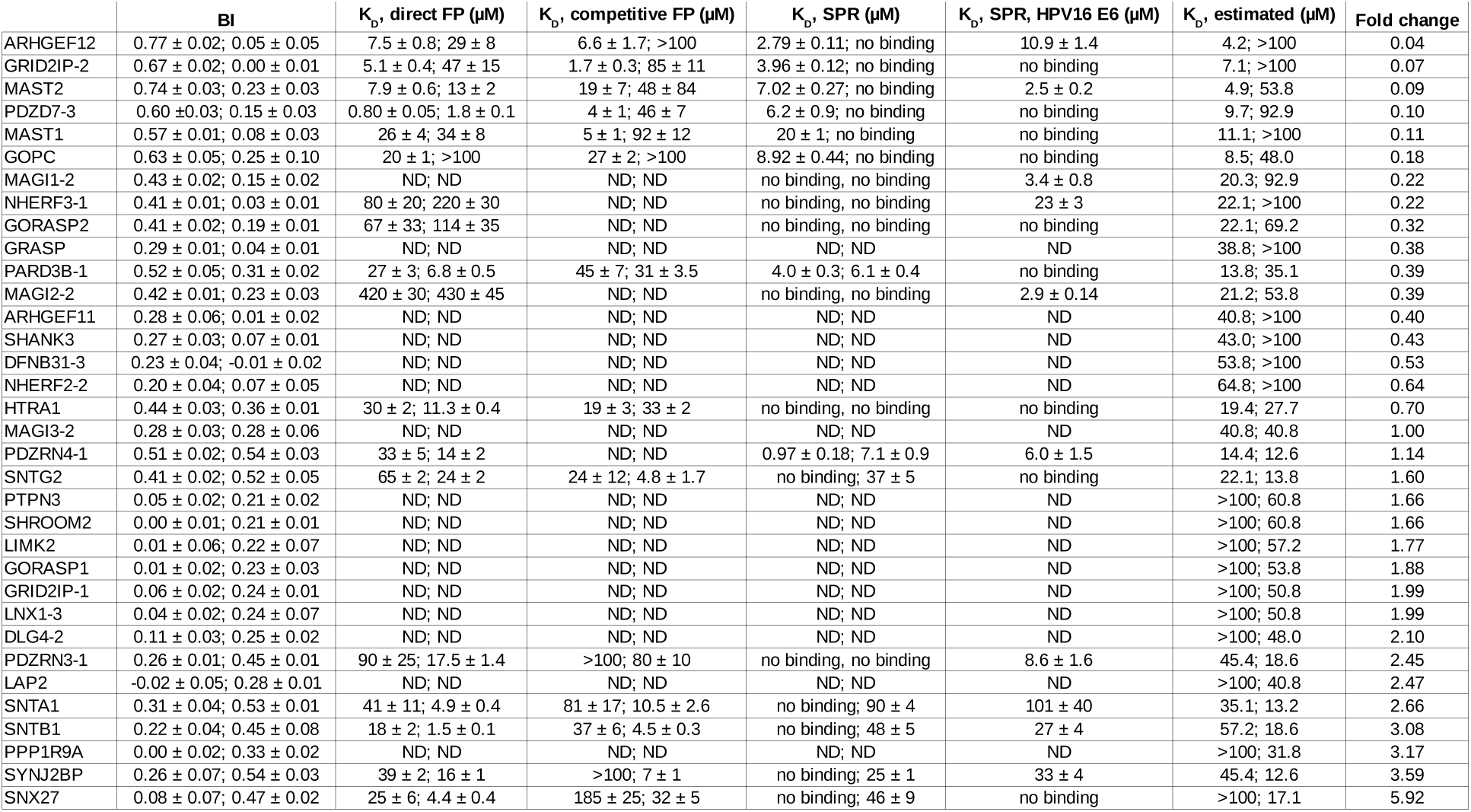
Summary of the *in vitro* validations. Values after the semicolon correspond to the phosphorylated RSK1 peptides. HPV16 E6 was used as an internal standard during the SPR measurements. K_d_ estimation was calculated from BI values as described in materials and methods and using an estimated 17 µM peptide concentration. Fold changes were calculated by dividing the estimated unphosphorylated and the phosphorylated dissociation constants. For undetectable interactions, a very weak K_d_ was assumed (100 µM, which corresponds to a BI of 0.14). ND means not determined, while no binding means that it was impossible to quantitatively measure their affinities in our experimental conditions.

### *In vitro* validation of PDZ-RSK interactions by biophysical approaches

To validate the holdup assay, we used orthogonal in vitro approaches. As a low-throughput yet unbiased validation, we measured the binding between long, unlabeled RSK1 peptides (683-735) and the PDZ domains of ARHGEF12 and SYNJ2BP in solution by ITC (Figure S2). These experiments confirmed that phosphorylation significantly increases binding to SYNJ2BP and decreases the affinity to ARHGEF12. Then, we decided to systematically test the interactions that showed a BI value larger than 0.4 in any of the two holdup assays. We investigated these interactions by SPR-BIAcore, as well as direct and competitive FP measurements (Figure 3, Table 2 and Figures S3, S4). With these techniques, we were able to accurately measure binding constants of 15, 36 and 28 PDZ-PBM pairs, respectively.

**Table 2.** Summary of the kinetic measurements. Dissociation rates were deduced from stopped flow fluorescence polarization experiments. On-rates are calculated based on the steady state affinities of the fluorescent peptides (deduced from direct FP measurements). The bias factor (the ratio of the binding affinities of the direct FP and the holdup assay) is the applied correction factor of the fitted dissociation rates to estimate the unbiased off-rates. The first half of the columns contain the parameters of the unphosphorylated peptide and the second half contain the parameters of the phosphorylated peptide. ND means not determined.

| | Measured $k_{off}$, fRSK1 ($s^{-1}$) | Calculated $k_{on}$, fRSK1 ($s^{-1}\mu\text{M}^{-1}$) | Bias factor, fRSK1 | Estimated $k_{off}$, RSK1 ($s^{-1}$) | Measured $k_{off}$, fpRSK1 ($s^{-1}$) | Calculated $k_{on}$, fpRSK1 ($s^{-1}\mu\text{M}^{-1}$) | Bias factor, fpRSK1 | Estimated $k_{off}$, pRSK1 ($s^{-1}$) |
| --- | --- | --- | --- | --- | --- | --- | --- | --- |
| ARHGEF12 | 46 ± 2 | 6 | 0.56 | 26 | ND | ND | 3.45 | ND |
| MAST1 | 129 ± 12 | 5 | 0.43 | 55 | ND | ND | 2.94 | ND |
| MAST2 | 101 ± 5 | 13 | 0.62 | 63 | ND | ND | 4.14 | ND |
| GOPC | 195 ± 18 | 10 | 0.43 | 83 | ND | ND | 0.48 | ND |
| GRID2IP-2 | 109 ± 5 | 21 | 1.39 | 152 | ND | ND | 2.13 | ND |
| PDZD7-3 | 72 ± 4 | 90 | 12.13 | 873 | ND | ND | 51.61 | ND |
| SNTG2 | ND | ND | 0.34 | ND | 364 ± 87 | 15 | 0.58 | 209 |
| SYNJ2BP | ND | ND | 1.16 | ND | 472 ± 67 | 29 | 0.79 | 372 |
| SNTA1 | ND | ND | 0.86 | ND | 610 ± 138 | 124 | 2.69 | 1643 |
| SNX27 | ND | ND | 4.00 | ND | 562 ± 85 | 128 | 3.89 | 2184 |
| PARD3B-1 | 676 ± 219 | 25 | 0.51 | 346 | 431 ± 48 | 63 | 5.16 | 2225 |

We used these datasets to estimate the correlation between measured BIs and the dissociation constants using Monte Carlo modeling and a general equation of the dissociation constant. Our calculations showed that the peptide concentration in the holdup assays should be around 14 and 23 µM (Figure 3A, Table 1). Using this fitted parameter, it can be calculated that the HU assay is capable of efficiently detecting any interactions with K_d_ < 65 µM (at the 0.2 BI cutoff).

Using the holdup assay, we have the unique opportunity to gain insight into the heterogeneous dynamic changes that occur after a single phosphorylation event (Figure 3B). At the two extremes, the phosphorylation results, according to the sensitivity thresholds of our binding assays, in OFF and ON switches, between RSK1 and ARHGEF12 or SNX27, respectively, but instead of detecting similar definite classes of switches, we have found that there is a real continuum between such effects. It is also clear that the amplitude of OFF signals is larger than the amplitude of ON signals, confirming the general notion that it is easier to destroy than to create. This continuum nicely reflects the gradual behavior of the system. Importantly, we detected significant interaction partners with either switch or dimmer and either ON or OFF functions. This suggests that there is no evolutionary pressure towards any kind of specific effects and that the system does not prefer any particular output (e.g. OFF switch or ON dimmer) (Figure 3B). In conclusion, we provided *in vitro* experimental evidence that phosphorylation reshuffles the whole RSK1-PDZ interactome.

### Dynamic rearrangement of the RSK1-PDZ interactome inside cells

The observed changes in steady-state binding affinities suggest large scale rewiring of the RSK-PDZ interactions. To test this concept, we validated selected interactions in a cellular context using a next generation split-luciferase fragment complementation system, called NanoBiT. This method is appropriate for detecting dynamic changes in PPIs (Dixon *et al*, 2016). Instead of using isolated PDZ domains and RSK peptides, we used full-length proteins in our assays in HEK293T cells. The N-terminus of RSK1 was tagged with the short NanoBiT tag (SmBiT) and either the N-or the C-terminus of the interaction partner with the large NanoBiT tag (LgBiT). Wild type (WT) and two mutant versions of RSK1 were used. The L714E mutation eliminates the interaction between ERK and RSK, therefore, it creates a constitutively inactive kinase (Alexa *et al*, 2015). In the case of the DC1 truncation, the last residue of RSK1 was removed, therefore, we eliminated the functional PBM of the protein (Gógl *et al*, 2018). To validate our interaction sensors, we compared the steady-state luminescence signals of these variations in serum starved cells. We measured high luminescence signals with the ARHGEF12, GOPC, PARD3B, MAGI1 and SYNJ2BP sensors (Figure 4A). The C-terminal truncation significantly reduced the luminescent signal in all cases, while the L714E mutation decreased the luminescence outputs in cases of PARD3B and SYNJ2BP. Note that in our *in vitro* experiments, the PDZ domains of these two partners interacted with the phosphorylated RSK1 PBM with a higher affinity, compared to the other monitored interaction partners. Thus, we were able to validate interactions between full-length proteins and RSK1, in a cellular context.

**Figure 4.**
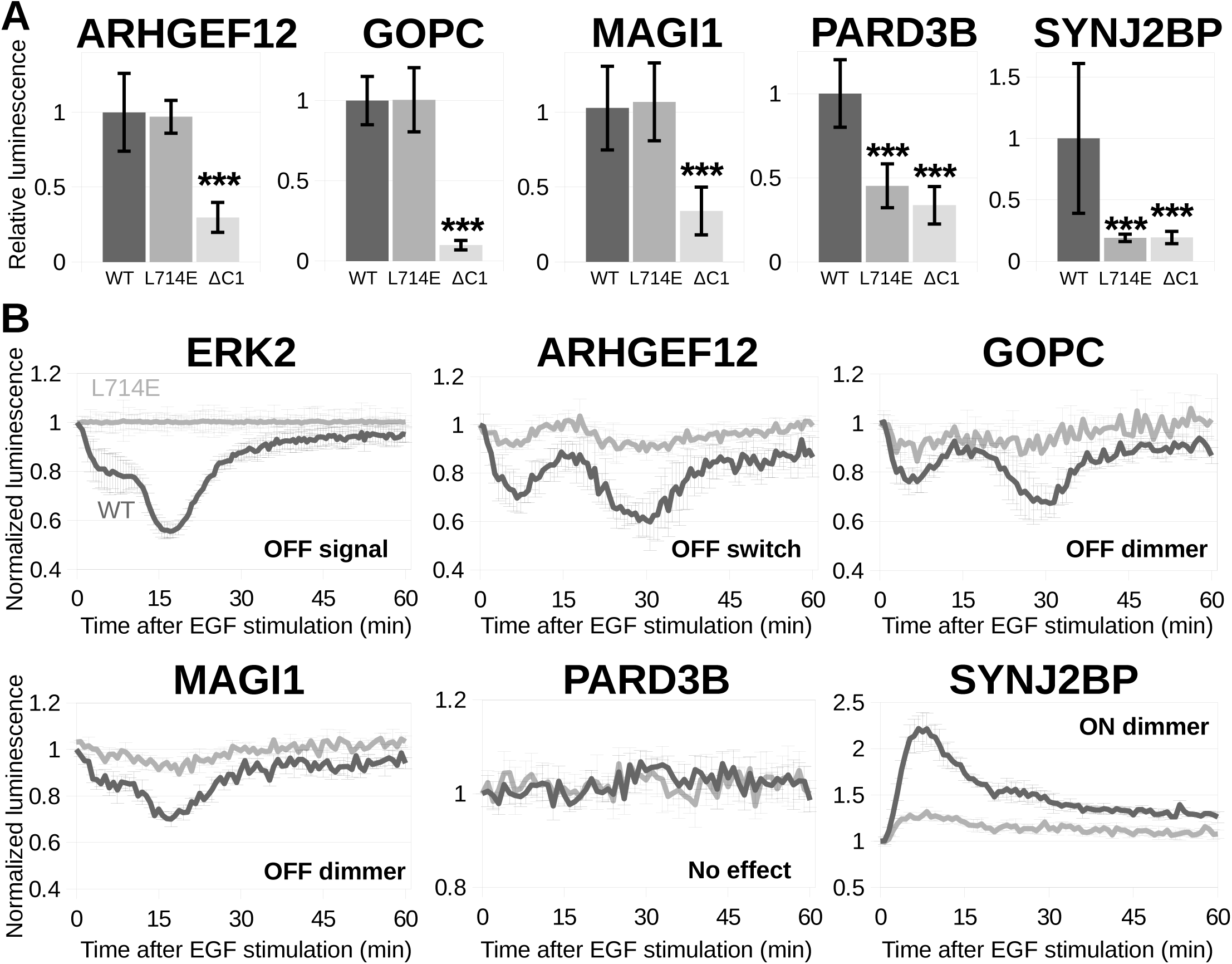
Live-cell monitoring of RSK1 binding to PDZ-containing partners and of its modulation upon EGF simulation. (A) Monitoring steady-state luminescence, with the interaction sensors between RSK1 and full length PDZ proteins. Full-length proteins fused to two complementary fragments of nanoluc luciferase were co-expressed in serum starved HEK293T cells. The resulting luminescence signal was measured as indicated in material and methods. The luminescence signal obtained for the pair of wild-type constructs is used as reference (relative luminescence). The L714E RSK1 mutant is known to eliminate the interaction between RSK1 and ERKs (Alexa *et al*, 2015). The ΔC1 RSK1 mutant does not contain the last C-terminal residue of RSK1 and therefore does not contain a functional PBM. The luminescence signal is systematically disrupted by the ΔC1 mutation, indicating that this signal efficiently reports the PBM-mediated binding of RSK1 to its PDZ-containing targets. The L714E mutation disrupts the signal in cases where the interaction partner can significantly interact with the phosphorylated form of RSK1. (n=6) Asterisks indicate statistical significance (*** P<0.001) calculated by two-tailed Student’s t-test between the luminescence signals of mutant and WT RSK1 constructs. (B) The steady-state validated RSK1 based luminescence interaction sensors (with ERK2 and several proteins containing RSK1-binding PDZ domains) were co-expressed in serum-starved HEK293T cells. The luminescence signal in absence and in presence of EGF (20 ng/ml) was monitored for sixty minutes following EGF addition. The measured luminescence signal was normalized to the initial luminescence and to the spontaneous substrate (furimazine) decay based on the unstimulated cells. The dark and grey curves show the luminescence signals of the WT and the L714E mutant, respectively. EGF stimulation provokes a time-modulated decrease of the luminescence signal for co-expressed constructs of RSK1 and ERK2 as observed in our previous work (Gógl *et al*, 2018). Note that EGF simulation diversely modulates (increase, decrease or no significant change) the luminescence signal for each PDZ-containing protein in a similar timescale of the RSK-ERK dissociation. Remarkably, this EGF-induced luminescence signal modulation in the cellular assay follows the same trend as the phosphorylation-induced modulation (off and on dimmer) of the *in vitro* binding affinity of RSK1 to individual PDZ domains.

MAPK pathway, RSK activation and its PBM phosphorylation, can be triggered in cell cultures by various stimulations (Cargnello & Roux, 2011). We used EGF stimulation in our NanoBiT assays to measure the changes in the luminescence signal. Extracellular-stimulation induced changes in the sensor brightness within the same timescale as of the ERK-RSK dissociation profile (Figure 4B). In all cases, the maximum change was detectable between 5-15 min and the signal started to disappear after 30-45 min. We observed periodic signals, as in our previous study, which seem to be a characteristic feature of RSK interactions (Gógl *et al*, 2018). ARHGEF12, GOPC and MAGI1 showed a decrease in luminescence after stimulation. In contrast to these OFF signals, PARD3B did not show any change after activation of the pathway and SYNJ2BP showed an increased luminescence after EGF stimulus. These observed trends were correlated with our in vitro measurements.

### A compendium of potential RSK targets

If a protein physically interacts with a protein kinase, it may also be its substrate. However, initial analysis of phosphoproteomic data revealed that many of the strong RSK1 PBM binders (like GRID2IP, GOPC, PDZD7 or PDZRN4) do not contain a single phosphorylation site matching the RSK1 consensus motif (Figure 2B) (Hornbeck *et al*, 2015). Therefore, we decided to take a more focused look on all of these proteins, as potential RSK substrates. For this reason, we collected RSK-focused phosphoproteomic datasets for a meta-analysis. To our knowledge, there are three such datasets. (i) Galan et al. searched for RSK substrates using specific inhibitors (Galan *et al*, 2014). (ii) Moritz et al. tried to find tyrosine kinase activated AGC kinase substrates (Moritz *et al*, 2010). (iii) Avey et al. used the viral ORF45 protein to activate the ERK-RSK axis in cells and they searched up-or down-regulated phosphoproteins (Avey *et al*, 2015). (iv) In addition, the ERK compendium has recently been published (Ünal *et al*, 2017). It is a systematic collection of ERK related phosphoproteomic studies containing both direct and indirect ERK substrates. The compendium is also a great resource for potential RSK phosphosites (RxxS and RxRxxS motifs) (Figure S5AB) (Galan *et al*, 2014). We used this subset of the ERK compendium as an additional resource to our meta-analysis. The four potential RSK substrate collection, termed here as RSK compendium, included 997 potential substrates, where 349 substrates were identified in more than one study (Figure 5A, Table S2). Only 35 substrates were identified in all of the phosphoproteomic datasets but this short list included some well characterized RSK substrates, such as ARHGEF12, EIF4B, EphA2, GSK3B, PFKFB2, PPP1R12A (MYPT1), RPS6, or SLC9A1 (NHE1) (Shi *et al*, 2018) (Shahbazian *et al*, 2006) (Zhou *et al*, 2015) (Lara *et al*, 2013) (Houles *et al*, 2018) (Artamonov *et al*, 2013) (Cuello *et al*, 2006).

**Figure 5.**
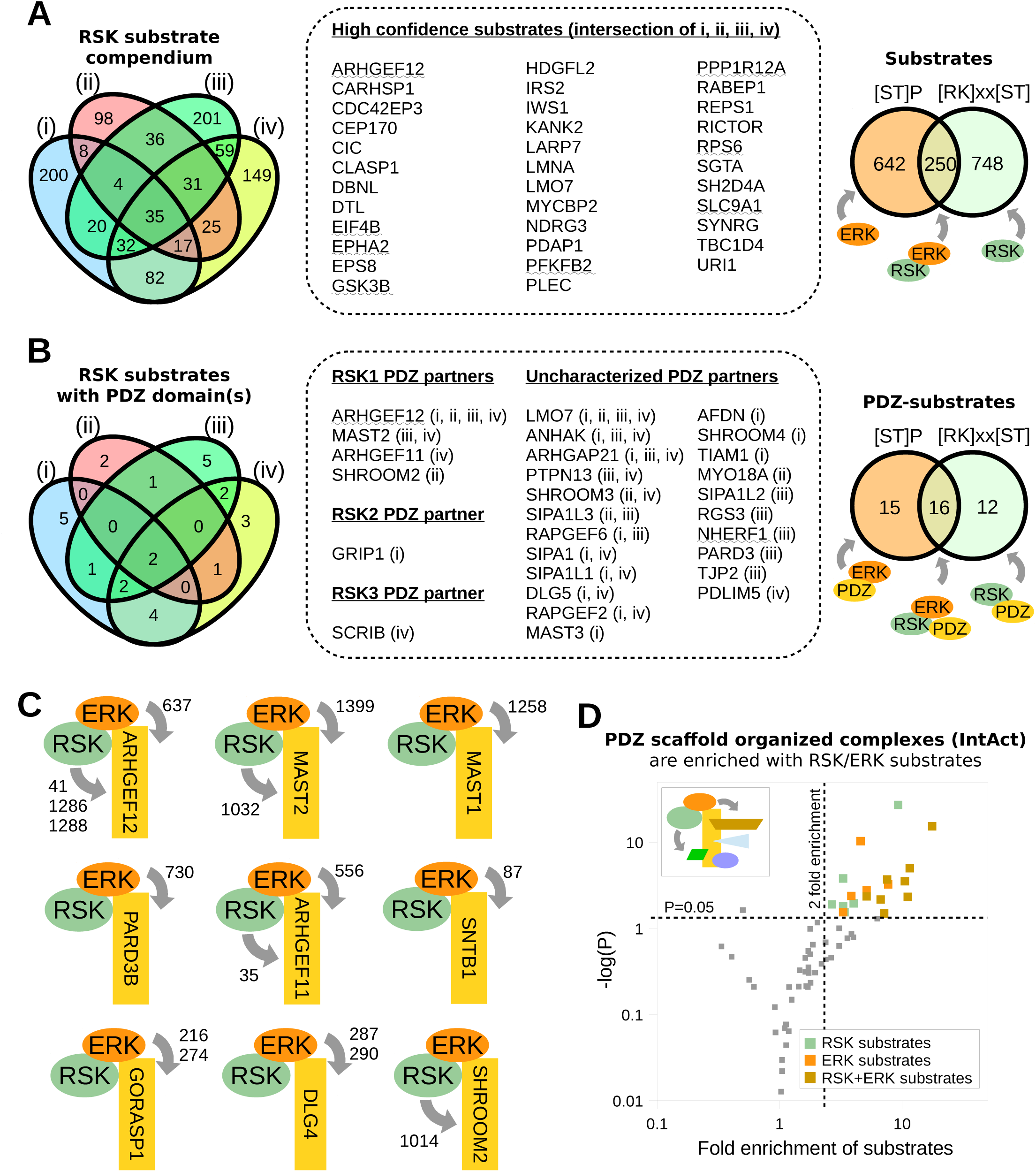
Meta-analysis of phosphoproteomic studies and bioinformatic search to find potential direct and indirect PDZ-dependent substrates of RSK and of ERK. (A) Left panel: a graphical representation of the intersections of RSK substrate lists from four different HTP phosphoproteomic studies: (i) Galan et al. (Galan *et al*, 2014), (ii) Moritz et al. (Moritz *et al*, 2011), (iii) Avey et al. (Avey *et al*, 2015), (iv) [RK]xx[ST] subset of the ERK compendium (Ünal *et al*, 2017). Middle panel: the intersection of the four lists contains several previously characterized RSK substrates (underlined), suggesting that other proteins found in this core intersecting ensemble may also represent high-confidence RSK substrates. Right panel:the RSK compendium and the direct ERK compendium significantly overlaps. This indicates that a set of substrates can be phosphorylated on both ERK ([ST]P) and RSK ([RK]xx[ST]) consensus sites. (B) Same representation as in (A), focusing on RSK substrates with PDZ domains. Only a few PDZ domain containing substrates are present in the whole dataset. Moreover, only ARHGEF12 was found in the core intersecting ensemble, and only a handful of PDZ interaction partners were found in more than one HTP study. Uncharacterized PDZ partners could be direct partners of other RSK isoforms, PDZ-independent substrates or false positives. (C) Many RSK1 PDZ interaction partners contain an ERK phosphorylation site. Additionally, a few substrates, such as ARHGEF12, can be phosphorylated by both kinases. (D) The IntAct database was used to estimate the enrichment of ERK and RSK substrates among the interaction partners of the RSK1 PDZ-dependent interaction partners. On the vulcano plot, each dot represents the enrichment of kinase substrates among the interaction partners of a PDZ scaffold. We have identified a high number of potential indirect RSK and ERK substrates among these interaction partners, which are indicated with colors in the upper right corner. P values indicate statistical significance compared to a random pool of intracellular proteins, calculated by Chi-square test. Fold enrichment indicates the increased proportions of substrates compared to the same random pool.

### Direct and indirect phosphorylation through scaffolding mechanisms by ERK and RSK

Only 28 potential RSK substrates were identified, which contained at least a single PDZ domain and most of these substrates were identified in a single dataset (Figure 5B). Moreover, most of these were not detected by the holdup assay as interaction partners of the RSK1 peptides. The most robust PDZ substrate of RSK is definitely ARHGEF12, the strongest *in vitro* interaction partner of the unphosphorylated RSK1 peptide, which was identified in all phosphoproteomic datasets. Additionally, MAST2, ARHGEF11 and SHROOM2 were also identified in our RSK substrate compendium. We have also found PDZ-dependent interaction partners of other RSK isoforms (Sundell *et al*, 2018) (Thomas *et al*, 2005) (Lim & Jou, 2016). This analysis revealed that there are only four direct RSK substrates among its potential 34 PDZ partners.

The RSK and the ERK compendiums show an overlap, therefore some substrates can be phosphorylated by both RSK and ERK (Figure 5AB). Although the MAPK-and the PDZ-binding motifs can be found in the same C-terminal tail region of RSK, separated by only a few residue linker, it is stereochemically possible to form a ternary complex between the three domains (Figure S5) (Gógl *et al*, 2018). Therefore, ERK can also phosphorylate RSK-bound PDZ proteins. We have found that many RSK1 interaction partners can be phosphorylated by ERK. For example, ARHGEF12 contains three RSK phosphorylation sites and a single MAPK phosphosite (Figure 5C). Aside from this binder, ERK can phosphorylate seven other RSK interaction partners. In conclusion, while RSK1 interaction partners are not perfect RSK substrates, some of them can be phosphorylated by ERK. In this situation, the C-terminal tail of RSK will serve as a scaffolding module, bringing ERK and PDZ substrates close to each other.

To identify additional indirect, PDZ-scaffold mediated substrates, all potential interaction partners of our RSK1 binding PDZ-scaffolds were collected from the IntAct PPI database (Kerrien *et al*, 2012). Although the coverage of mammalian interactomes is still limited and these databases contain a high background, we observed significant enrichment of RSK and ERK substrates in many cases. The most promising scaffold was MAGI1, which was not identified previously as a direct substrate of RSK (or ERK). MAGI1 has a decent interactomic coverage (with 74 interaction partners) and more than 40% of these interaction partners were found to be potential RSK substrates. Similarly, 30% of the interaction partners are potential ERK substrates and 18% of them can be phosphorylated by both RSK and ERK (Figure 6D, Table S3). We have found similarly significant enrichment of RSK/ERK substrates among various interaction partners, such as ARHGEF11. In conclusion, while we have found that only a small portion of RSK1 binding PDZ proteins are direct substrates of RSK1, we have strong indications that many of them are scaffolds and there are many relevant potential RSK and ERK substrates among their interaction partners.

**Figure 6.**
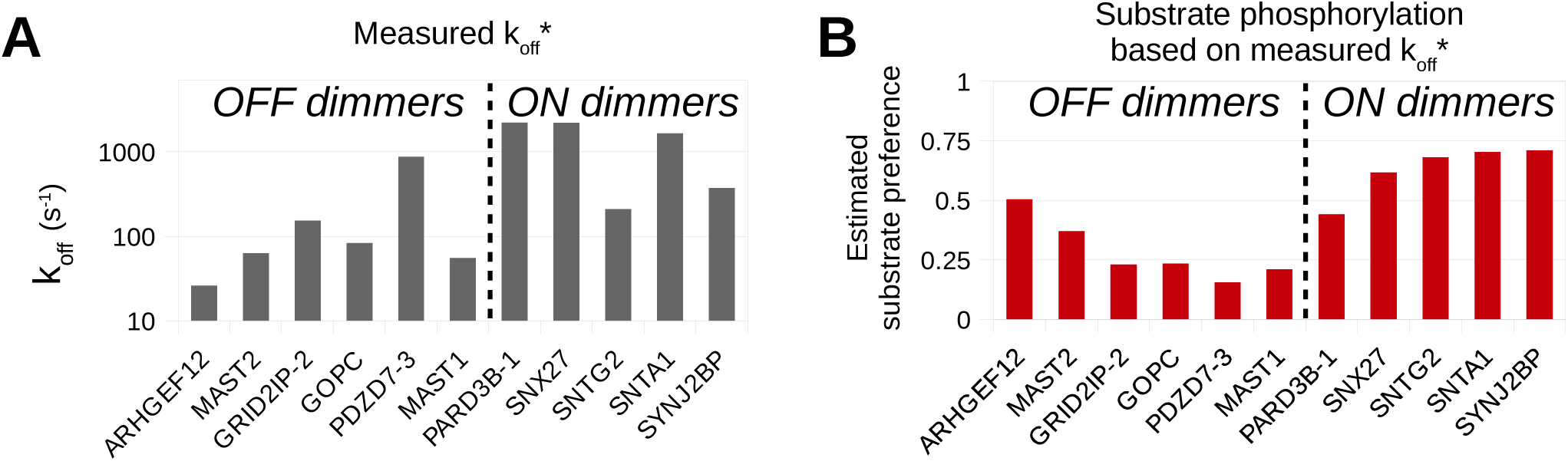
Binding kinetics differ between ON and OFF dimmers. (A) Dissociation rates were measured by the direct fluorescence polarization changes in a stopped-flow setup for a set of RSK1-PDZ interactions. Partners with OFF dimmer behavior showed a slower binding kinetics while the ON dimmers preferred faster binding rates. (B) Substrate phosphorylation was *in silico* estimated using their measured dissociation rate. Asterisks denote that the measured dissociation rates were already corrected to unbiased binding of unlabeled peptide.

### Kinetic control of feedback coupled substrate phosphorylation

To measure kinetic parameters, especially dissociation rates, we used a stopped-flow instrument, equipped with a home-made polarization toolkit (Figure S6). We rapidly mixed PDZ saturated fluorescent peptides with high molar excess of unlabeled peptides and continuously monitored the change in fluorescence polarization. This way, we can measure the dissociation rate of the fluorescent peptides (Table 2). Although the fluorescein labeling altered the steady-state affinity of some interactions (Table 1), we could assume that it only affected the dissociation rates as it is a large hydrophobic group. Therefore, we can estimate the unbiased off-rates by a correction to the differences in the steady-state affinities (of labeled and unlabeled peptides). Our results revealed that OFF dimmers have a generally slow binding kinetics (average k_off_ ≈ 230 s^-1^) while ON dimmers showed faster dissociation rates (average k_off_ ≈ 1300 s^-1^) (Figure 6). We used an *in silico* network-based modeling software to estimate substrate phosphorylation efficiency using these obtained kinetic parameters (Figure S7) (Harris *et al*, 2016). The amount of phosphorylated substrate was estimated in these simulations, induced by the same amount of external stimulation, assuming strong ON or OFF behaviors. The simulation demonstrated that the slower dissociation rate of OFF dimmers compensated their low substrate phosphorylation efficiency.

### Refining the role of RSK in RhoA activation

The RSK1 phophorylation site on ARHGEF12 lies next to a conserved motif, which was previously annotated as a putative CRM1-dependent nuclear export signal (NES) or a NES enhancer (Grabocka & Wedegaertner, 2007). To test whether the RSK phosphorylation alters ARHGEF12 localization, we transfected HEK293T cells with mCherry-fused WT and with S1288E phosphomimicked ARHGEF12 constructs (Figure S8). We have found that ARHGEF12 showed a clear cytoplasmic localization, despite the presence of the phosphomimetic mutation. Therefore, this phosphorylation site and the uncharacterized motif is probably not a NES-switch. Future studies should reveal the real physiological role of this putative phosphorylation regulated linear motif.

Despite the lack of changes in the protein localization, previous experiments showed that RSK phosphorylation affected the GEF activity of ARHGEF12. To gain insights into these mechanisms, we first measured the effects of MEK and RSK inhibitors on RhoA activation of ARHGEF12 overexpressing cells (Figure 7A). We have found that exposure to a MEK inhibitor decreased, and a RSK inhibitor increased the GTP bound RhoA levels in cells. Similar result was observed by using an overexpressed phosphomimetic mutant ARHGEF12 (Figure 7B). We transfected the cells with the wild type, the S1288E phosphomimetic and a W769D mutant version of ARHGEF12. The W769D mutation is near the RhoA binding site of the DH-PH domains, and *in vitro* it decreased the RhoA binding sevenfold (Kristelly *et al*, 2004). Inside cells, the W769D mutant decreased the overexpression-induced effect of ARHGEF12, it induced significantly smaller amount of active RhoA. Interestingly, the S1288E phosphomimetic mutant also decreased the signal by 20%. This resembles our previous experiment, where RSK inhibition increased the GEF activity. In conclusion, the RSK1 phosphorylation site lies next to a functionally important site that negatively affects the activity of ARHGEF12 under sustained conditions.

**Figure 7.**
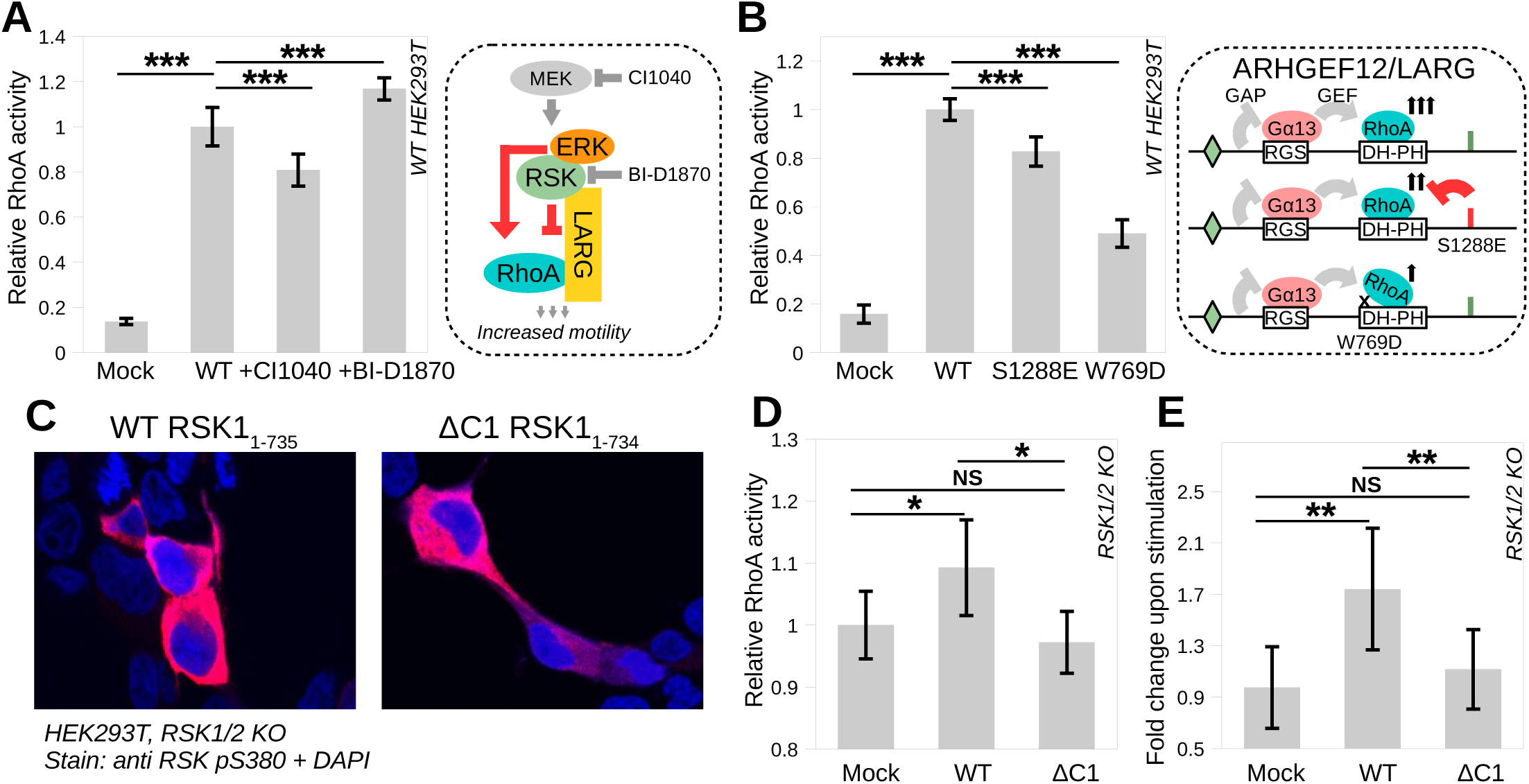
A possible RSK1-mediated cross talk between the ERK and the RhoA pathways. (A) Inhibition of the MAPK pathway alters levels of GTP bound RhoA. To visualize this effect, we overexressed WT ARHGEF12 in HEK293T cells, which resulted in a significant increment in the basal active RhoA levels. While MEK inhibition decreased, RSK inhibition increased the intracellular active RhoA level. (n=4) (B) Mimicking the RSK1 phosphorylation site on ARHGEF12 (S1288E) or introducing a RhoA binding incompetent mutant (W769D) altered the RhoA signalization. Phosphomimicking decreased the signal by 20% and W769D mutation by 50%. (n=4) The schematic model of ARHGEF12/LARG activation is highlighted on the right side, including GAP and GEF activities. (C) RSK1/2 knockout HEK293 cell line was used to measure the role of the PBM of RSK1 in a more native environment. Deletion of the RSK1 PBM does not affect the localization of active RSK1. (D) The presence of intact RSK1 increases the basal RhoA activity but this effect is decreased without a functional PBM. (n=4) (E) Transfected and serum starved cells were stimulated with serum (20%, 5 min). Without intact RSK1 (in the mock transfected knockout cell or in the presence of the PBM-lacking RSK1 construct) only minor increment was observed in the RhoA activity. The presence of intact, wt RSK1 enabled a proper response in RhoA activation upon stimulation. (n=4) Asterisks indicate statistical significance (** P<0.005, * P<0.01, NS P>0.01) calculated by two-tailed Student’s t-test.

To reveal the importance of the functional PBM of RSK1 on RhoA activation, we used a double RSK1/2 knockout HEK293 cell line. This cell line helped us to investigate RSK phenotypes without the endogenous background. To test whether the PBM affects the localization of RSK1, we transiently transfected this cell line with either an untagged full-length (1-735), or a single residue truncated version (1-734) of RSK1 clone or with an empty vector. We have found that the overexpressed and phosphorylated RSK1 localized in the cytoplasm, similarly to the endogenous phospho-RSK in a wild type HEK293T cell line (Figure 7C) (Gógl *et al*, 2018). In contrast, an increased level of the basal RhoA activity was only apparent in the presence of the WT RSK1 construct (Figure 7D). This slight increase was more apparent in a serum stimulation dependent experiment. Here, the stimulation induced large change in RhoA activity in the presence of the WT RSK1 construct, but not with the mock or the shortened RSK1 protein (Figure 7E).

## DISCUSSION

### Regulation of RSK1-PDZ interactions by PBM autophosphorylation

Previously, only a handful of PDZ interaction partner of RSK1 had been identified and their response to PBM Ser732 autophosphorylation was largely unknown. Here, we attempted to fully describe the PDZ interactome of RSK1 including its regulation by PBM autophosphorylation at Ser732. Altogether, 34 interaction partners were identified, most of them being novel with the notable exception of MAGI1. By contrast, we did not detect any interaction between NHERF (EBP50) or the first PDZ domain of Scribble in contrast to previous reports about the PDZ-dependent interaction of these two proteins with RSK1 (Lim & Jou, 2016) (Sundell *et al*, 2018). Although most of our interactions were affected by PBM phosphorylation, we have found only a few cases that can be considered a genuine phospho-switch within the detection threshold of our approaches. For example, binding to ARHGEF12 and GRID2IP were mostly eliminated, while binding to the adapter protein SNX27 was promoted by phosphorylation. In contrast, most substrates showed a dimmer effect and approximately as much ON as OFF dimmers were identified. These partners are able to interact with both states of the peptide, but with different affinities. The rest of the interaction partners (such as PARD3B) mediated highly similar interactions with both states and therefore, these interaction partners are unable to sense the presence (or the absence) of a phosphate group.

Mitogenic stimulation activates the ERK pathway, such as EGF stimulation, which activates the EGFR pathway (an upstream activator of the ERK cascade). Eventually, the downstream signals will activate the MAPK (ERK), which leads to the phosphorylation of RSK. The Ser732 phosphorylation site is a well-defined major autophosphorylation site on RSK, therefore, upon stimulation, we can expect dynamic changes in the RSK PBM-PDZ interactome based on quantitative *in vitro* measurements and *in silico* simulations. To test this hypothesis, we created five intracellular PPI sensors for PDZ dependent RSK1 interactions. In the holdup assay, ARHGEF12, GOPC and MAGI1 showed a preference for the native PBM, while the PDZ domain of SYNJ2BP preferred the phosphorylated peptide. PARD3B slightly preferred the native peptide in the holdup assay, but our *in vitro* validations (competitive FP and SPR) revealed that the difference between the two affinities is marginal and therefore, it should interact with both versions of RSK1. The obtained changes in the brightness of the luminescent PPI sensors showed the same trends. The PPI sensors of ARHGEF12, GOPC and MAGI1, the OFF dimmer interactions, showed a drop in their luminescence signals. In contrast, on one hand PARD3B did not showed any change in the luminescence upon EGF stimulation, and on the other hand SYNJ2BP, an ON dimmer interaction, showed a large increment in luminescence after stimulation. In conclusion, we could show that our proposed phosphorylation-induced PDZ interactome rewiring occurs also inside cells, in an EGF stimulation dependent manner.

### PDZ interactions are important for RSK1 substrate targeting

Of all known phosphorylation sites in the PDZ binding partners of RSK1, we noticed that many of them lack potential RSK phosphorylation sites, suggesting that these interaction partners are rather scaffold proteins than direct substrates. However, there were several exceptions, such as ARHGEF12 and MAST2. These partners contained many potential phosphorylation sites, matching the RSK consensus site. Moreover, some of them can be found in the RSK and ERK substrate compendiums. Out of the 34 interaction partners of RSK1, we identified 4 direct RSK substrates and 5 additional substrates which can be potentially phosphorylated by ERK in the ERK-RSK-PDZ ternary complex. This analysis suggests that the rest of the interaction are rather scaffold proteins than substrates. In this manner, they are also actively involved in substrate targeting, therefore the RSK1-PBM is an essential element in the function of these kinases.

Interestingly, ARHGEF12 and MAST2 phosphopeptides were found in the ORF45 phosphoproteomic dataset (iii). ORF45 is a viral protein of the Kaposi’s sarcoma-associated herpesvirus, which causes sustained RSK1 activation by stabilizing the ERK-RSK complex and by preventing their dephosphorylation (Kuang *et al*, 2009). As the viral protein activates RSK1, it likely causes constitutive RSK-PBM phosphorylation. Both of these interactions were characterized as strong OFF dimmers in our experiments, their interaction should be minimized under viral infection. Indeed, the presence of the viral protein down-regulated the phosphorylation of both substrates.

Among the unambiguously identified RSK substrates, ARHGEF12 has a dedicated place. It is a strong partner of the RSK1 peptide and their interaction shows a dissociation profile upon EGF stimulation. Moreover, Shi et al. have recently showed that the association between RSK2 and ARHGEF12 (also known as Leukemia-associated RhoGEF or LARG) is essential in RhoA activation in glioblastoma cells (Shi *et al*, 2018). They discovered that RSK can interact with this RhoA GEF and it can indeed phosphorylate it at Ser1288. They demonstrated that the presence of RSK is essential for the association between RhoA and ARHGEF12, and therefore, RhoA activation.

ARHGEF12 also has a GAP activity towards the Gα subunits. Its RGS domain can interact with activated Gα12/13 subunits, which will enhance the RhoA GEF activity of the protein (Kristelly *et al*, 2004). Therefore, activation of ARHGEF12 is kinetically regulated by the interplay of negative and positive feedbacks, between the bound Gα, RhoA and other factors, such as the RSK phosphorylation site. This phosphosite is located in the disordered tail of the protein. The complete removal of the tail affects both the oligomerization state and the localization of ARHGEF12 (Grabocka & Wedegaertner, 2007). While this phosphorylation site is not ancient (it can be found in amniotes) it lies right next to a fundamentally conserved site, which was previously highlighted as a putative CRM1 dependent NES or NES enhancer. In our experiments, S1288E phosphomimication did not altered the localization of ARHGEF12, but it affected its overall GEF activity. The phosphomimetic mutation decreased the intracellular GTP bound RhoA levels. Similar effect was observed when the cells were incubated with MEK or RSK inhibitors. However, it was impossible to reveal the precise molecular function of this delicate system by measuring its activity after a long-term exposure to an inhibitor or using a constitutive phosphomimetic mutant. However, using a RSK knock-out cell line, we detected a RSK1-PBM-mediated activation of the RhoA pathway. This is in line with the observations of the Ramos group, who found that the homologous RSK2-ARHGEF12 interaction is important in (serum, EGF, TNFα and PMA) stimulation dependent RhoA activation. These observations contribute to the increasing evidence, that a cross-talk occurs between the RSK paralogs and RhoA activation (Larrea *et al*, 2009).

### The importance of time in a dynamic network

Our meta-analysis showed that some of the PDZ mediated interaction partners of RSK are substrates or they can be part of larger assemblies, which contain an indirect substrate. However, in the case of RSK, phosphorylation of an OFF switch (or a strong OFF dimmer) is an apparent (veridical) paradox. To understand this, we must take into account that the interaction occurs only with the unphosphorylated tail, but only the active kinase will be able to phosphorylate the target sites. However, the activation will involve autophosphorylation of (the unbound) tail, eliminating the possibility of the interaction itself. Since these are weak complexes, the unbound fraction is relatively large, and as the autophosphorylation is a cis regulatory event, therefore it should follow a very fast kinetic rate. As a consequence, the phosphorylation of these targets should be negligible or they should follow a single turnover mechanism. (Which should result in a minor phosphorylation thanks to the weak binding affinity of PBM-PDZ interactions.) However, we could unambiguously detect some of them (like ARHGEF12) in phosphoproteomic datasets.

An optimally slow dissociation rate (off-rate) of the PDZ domain binding can resolve this paradox. It should maximize the lifetime of the PDZ-PBM complex at the beginning of the activation cycle, leaving enough time for substrate phosphorylation but it should be fast enough for maximizing the substrate exchange at the same time. Conversely, a fast off-rate is preferred in the case of an ON dimmer. In this case, the enzyme should maximize the substrate turnover with fast binding kinetics. Indications for this kinetic compensation can be found in our fast-kinetic measurements, where OFF dimmer interactions showed slower dissociation rates compared to the fast dissociation rates of ON dimmer interactions (Figure 6). Additionally, our *in silico* model confirmed this thought experiment (Figure S7). In this model, time is a very important parameter as signals will trigger a proportional positive or negative feedback. The lifetime of a complex (which is proportional to the dissociation rate constant of the docking interaction) will essentially limit the robustness of the signal transduction (Corzo & Santamaria, 2006). Without this kinetic compensation, a negative feedback mechanism should cause a single turnover cycle, where each activation step should trigger the phosphorylation of a tiny amount of substrate.

Therefore, substrate phosphorylation should not be exclusive to ON dimmers and kinetic compensation can enhance any substrate phosphorylation rate (Figure 6). This hypothesis also demonstrate that simple measurements of steady-state binding affinities (without kinetic information) are not enough to predict the quality of a substrate. A nice example is ARHGEF12, a definite OFF switch, which is a known and robust direct substrate of RSKs and not surprisingly, it showed the slowest binding kinetics in our measurements (Figure S6). We should emphasize here that these are general principles and they should be true for many other feedback-mediated enzymatic reactions.

### PDZ specificity and phosphorylation-sensitive PDZ domains

In our experiments, ARHGEF12 was the strongest partner of the RSK1 peptide. Interestingly, its very close paralog, ARHGEF11 mediated a much weaker interaction with the same peptide. A similar observation was made with the RSK1-like C-terminal peptide of CXC chemokine receptor 2 (their last four residues are identical) (Larrea *et al*, 2009). In their PDZ array screen, the peptide mediated much tighter interaction with ARHGEF12 than with ARHGEF11. There are only a few sequence differences between the paralogs. Moreover, all of the changes are within external loops, far from the peptide binding site (Figure 8A). This suggests that the specificity of a PDZ domain could not be reliably predicted based on purely the binding site itself. Additional parameters need to be considered, such as domain flexibility and stability, to reveal the driving force of PDZ-PBM specificity.

**Figure 8.**
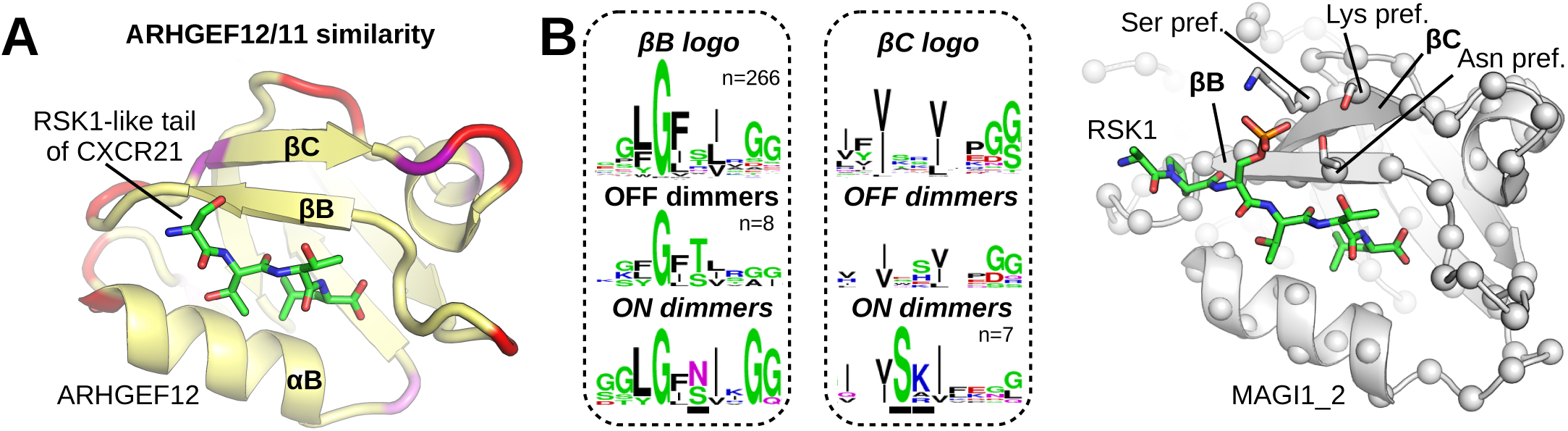
Determinants of PBM and -3 phospho-PBM specificity. (A) The close paralogs, ARHGEF12 and ARHGEF11, display different binding affinities with the unphosphorylated RSK1 peptide. Despite no alteration can be observed in their PBM binding groove, a handful of differences can be found in surface exposed loops. Different residues are colored in red and similar residues are colored in purple. Identical residues are colored in yellow. (B) A few differences can be observed in the PBM binding groove in ON dimmer PDZ domains. Sequence logos were generated from every human PDZ sequences or from identified dimmer subsets of the RSK1 peptide. Important differences are underlined in the ON dimmer sequence logo and their sidechains are showed with sticks in the structure of the OFF dimmer MAGI1 indicated with the preferred residues. In contrast, no preferences can be found for OFF dimmers.

Phosphorylation of PBMs is a very common regulation (Sundell *et al*, 2018). Based on our experiments, we identified a set of PDZ domains, which are responsible for mediating OFF or ON dimmer behavior of the -3 position site phosphorylation. Comparison of PDZ sequences reveals that there is no obvious driving force behind OFF dimmer behavior, but there are at least three positions within the peptide binding groove that can be important for ON dimmers (Figure 8B). The first of them is the outward facing residue of the second strand of the PDZ domain. This side chain is positioned in close proximity to the phosphate group and while it is usually a Ser/Thr residue in PDZ domains, an Asn residue is preferred within ON dimmers. The other two altered side chains are within the third strand of the PDZ domain. Here, both external side chains are altered in ON dimmers. Interestingly, the closest residue to the phosphosite is most frequently a Ser residue and the other one is a basic amino acid. Two of these residues are present in SNX27. Moreover, their interactions were already captured with -3 phospho-PBMs (Clairfeuille *et al*, 2016). Asn56 from βB and Ser82 from βC mediates a hydrogen bond with the phosphate group of PBMs. Moreover, replacing the basic residue in the βC (Arg762) to Ala in Scribble can swap the RSK3 binding properties from ON-to OFF-dimmer (Sundell *et al*, 2018). These observations lead us to the conclusion, that ON dimmer propensity is determined by the presence of phosphate acceptor sites and OFF dimmer propensity is probably relies on the PDZ specificity itself. We think that it is likely, that a very precise PDZ classification can be made, which will shows that a certain domain can interact with phospho-PBM (phosphorylated at a specific position) or not. (For example, the presence of the above residues are indications of a -3 phosphopeptide sensitive PDZ domain.) However, further studies are needed to collect more evidence about similar effects.

### Response to phosphorylation: switches and dimmers

Phosphorylation can alter linear motif binding by multiple ways. In the literature, most examples of phosphorylation induced PPI changes are considered as switches (usually called “phospho-switches”), which, can turn on, or turn off PPIs. However, many modulation is not binary and in some cases phosphorylation only slightly alters the binding itself, and it only acts as a simple fine-tuning mechanism. Taken out of context, such small effects can be considered as a switch but compared to a strong binary (“yes” or “no”) phosphorylation switch, which completely eliminates or triggers the binding, it is obvious that it is actually a minor effect. For this reason, we have recently proposed the term of ON/OFF dimmers in addition to ON/OFF switches. The term dimmer (or dimmer switch) is more appropriate to describe PPI rearrangements, as most phosphorylation-induced changes are gradual or stepwise and ON/OFF switch-like effects are likely to be only the extreme cases. Despite the obvious concerns regarding the switch like effects, the misleading “phospho-switch” term is generally used in the literature without using some softening expression, like the dimmer effect (“phospho-dimmer”). Our results demonstrate that there is a continuum between ON and OFF switches, including many gradually altered dimmers. Therefore, the dimming mechanism seems to be a more general description of the effect of phosphorylation events, while binary ON and OFF switches are extreme cases.

Considering all the above results and the number of players in the intracellular signaling pathways, it is tempting to claim that the dynamic rearrangement of the RSK1 interactome profile is just the tip of the iceberg. Recent technical advancements or future techniques will certainly allow us to shed more light on the full dynamics of the cell’s protein-protein interaction network.

## Supplement

**Figure S1.**
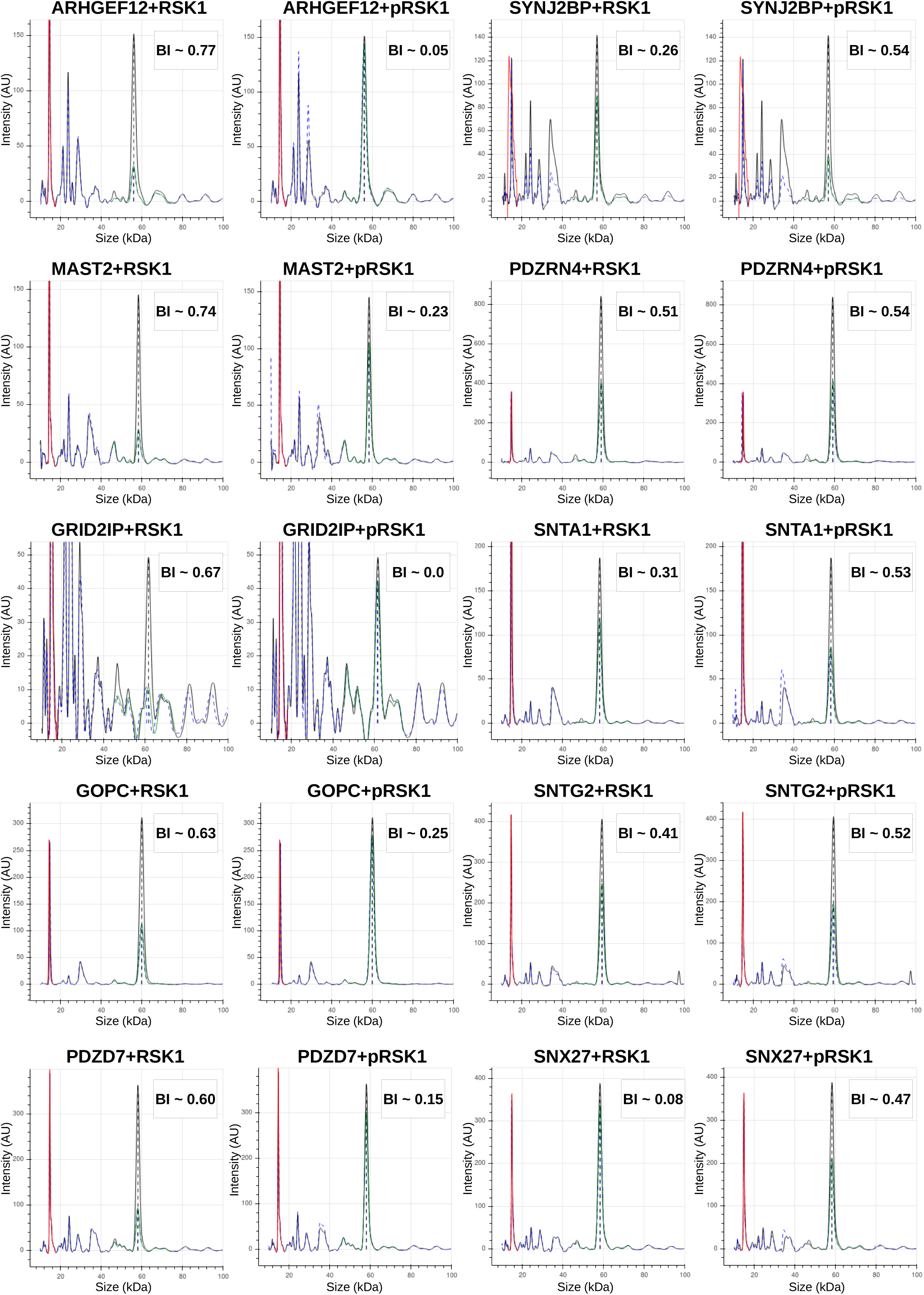
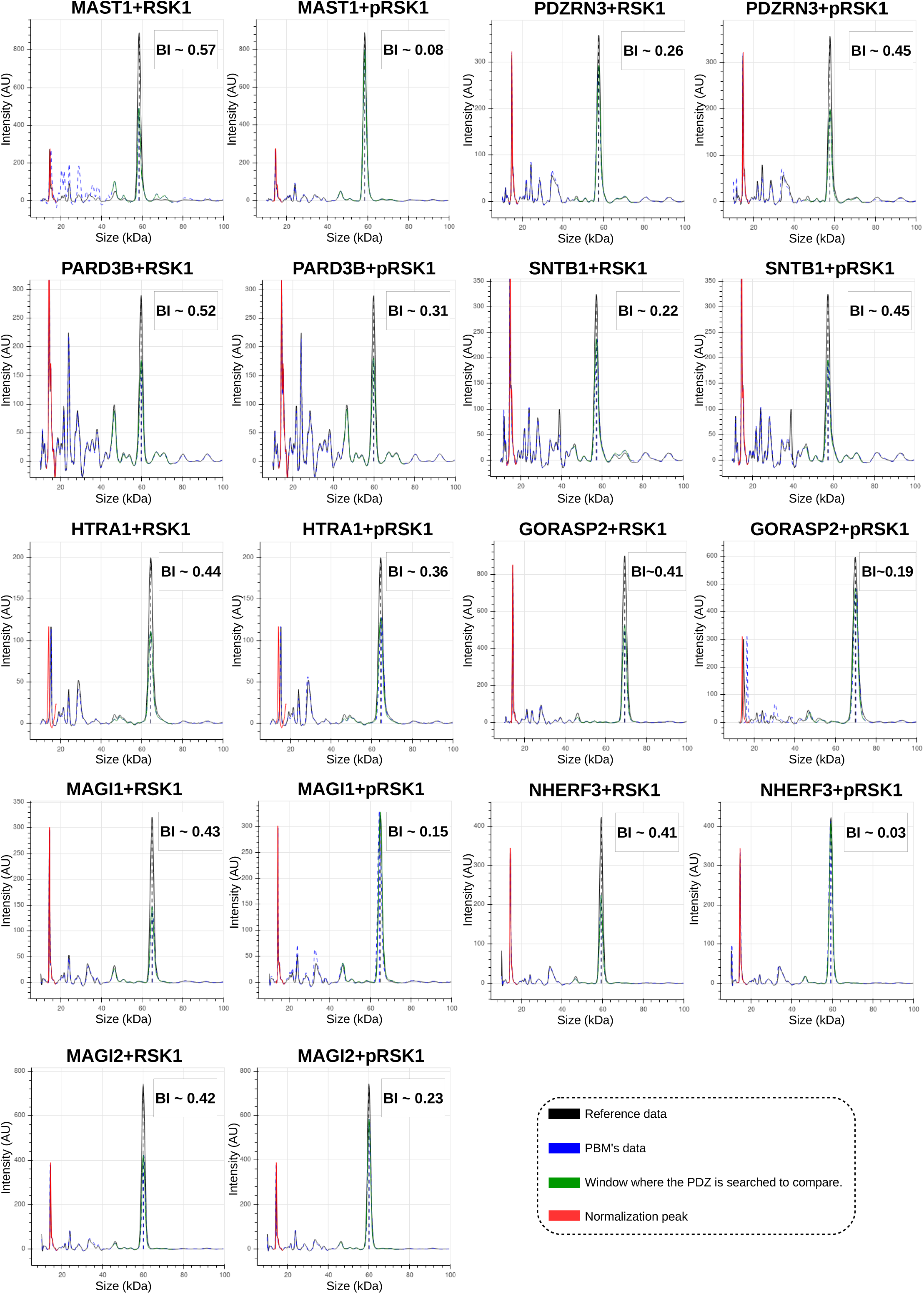
Raw results of the HU assay measured by the caliper. The overlay of processed electropherograms between the biotin control and the peptide experiment is shown for the most significant interaction partners of RSK1. The average BI value is highlighted in each panel. The more depleted the PDZ peak in the peptide experiment, the stronger the binding of the PDZ domain to the peptide.

**Figure S2.**
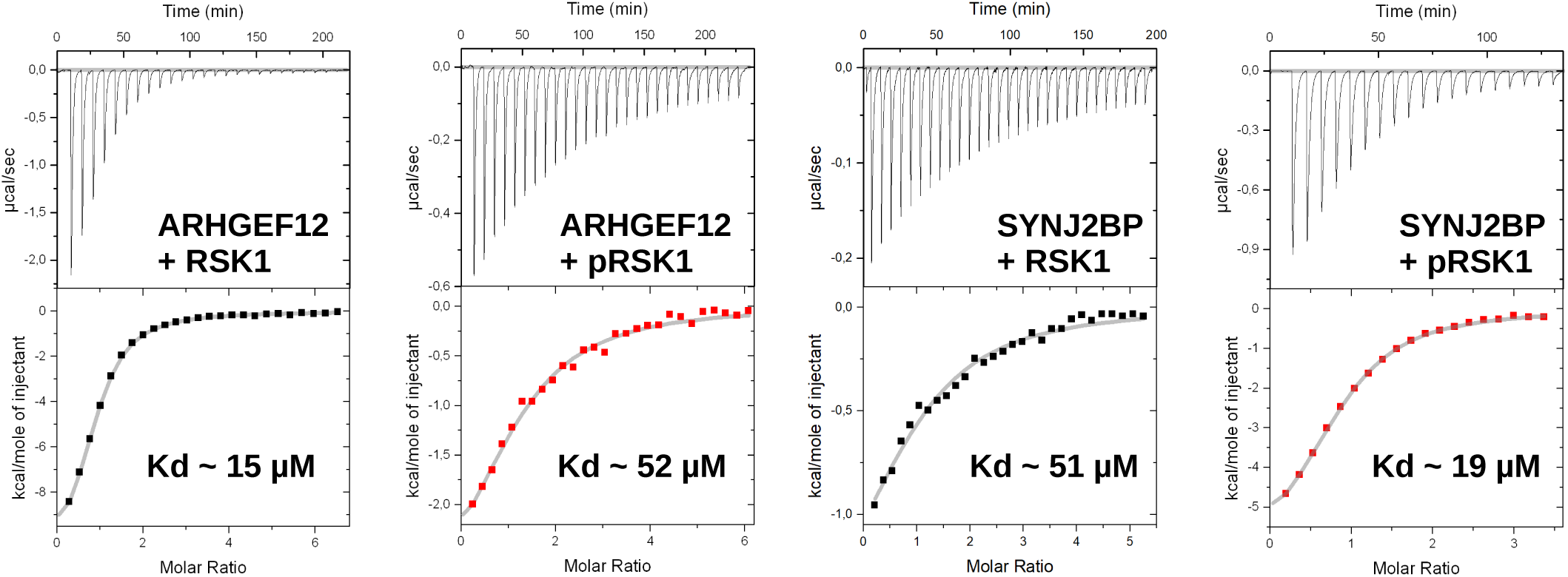
Results of the ITC measurements. Binding experiments were performed between the RSK1 and the PDZ domains of the strongest interaction partners of each peptides (of ARHGEF12 and SYNJ2BP) at 37°C. The calorimetric measurements confirmed the differential binding upon phosphorylation.

**Figure S3.**
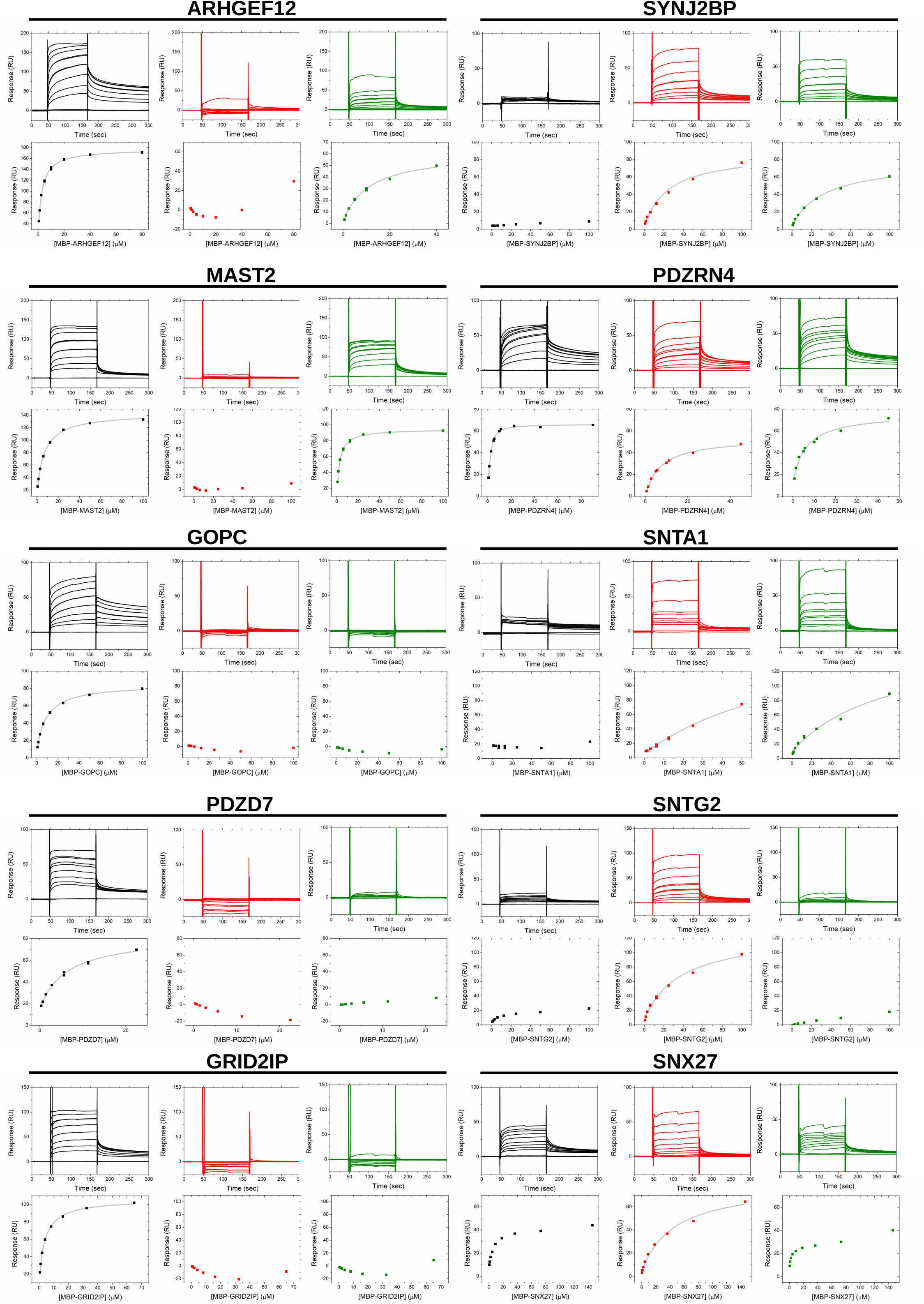
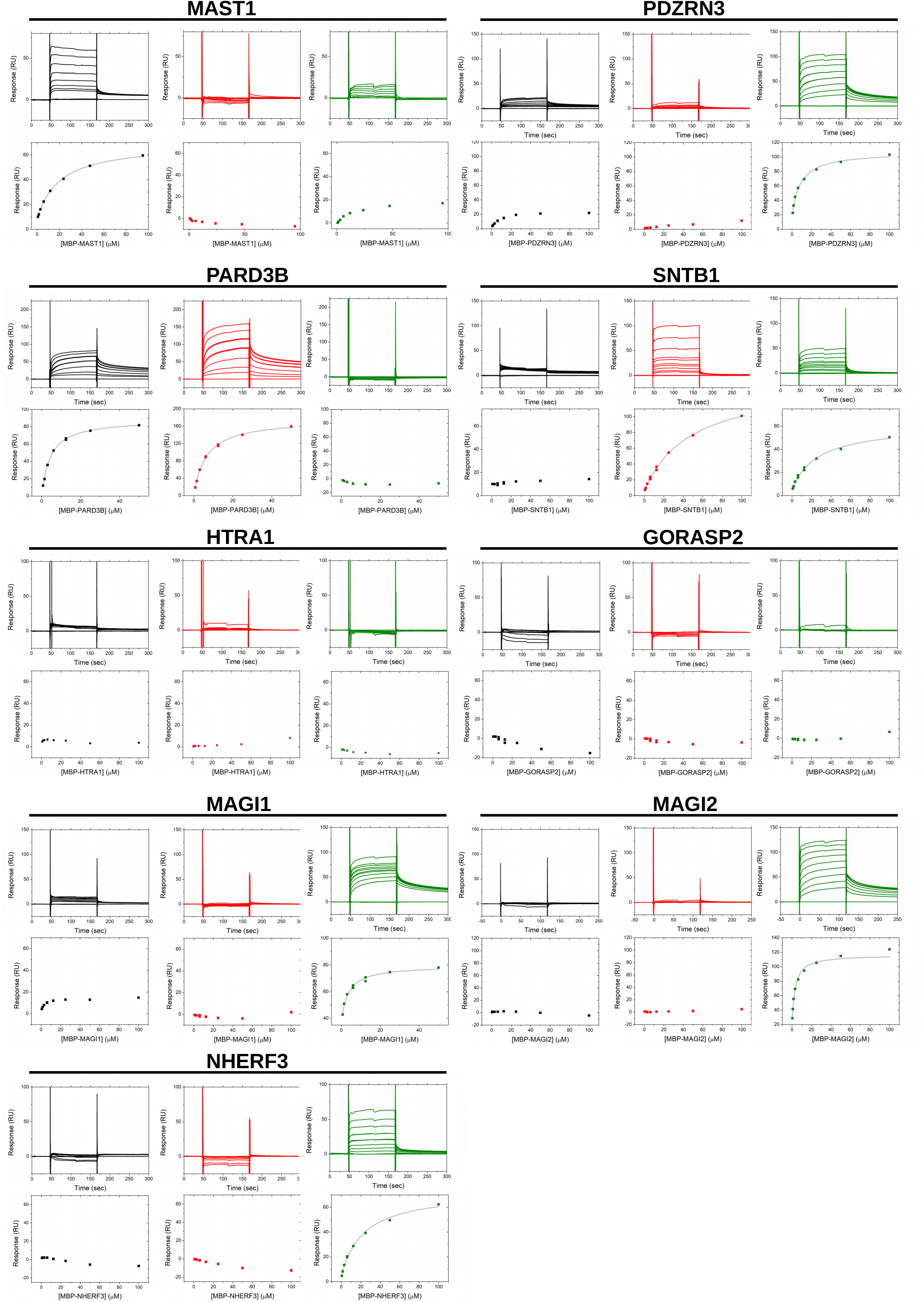
Results of the SPR measurements. The four channel of the CM5 chip was split into a negative control, a HPV16E6 internal control (green), an unphosphorylated (black) and a phosphorylated (red) surface and the significant RSK1 interaction PDZ domains (fused to MBP) were injected into the surface. Only steady state analysis was performed due to biphasic sensograms.

**Figure S4.**
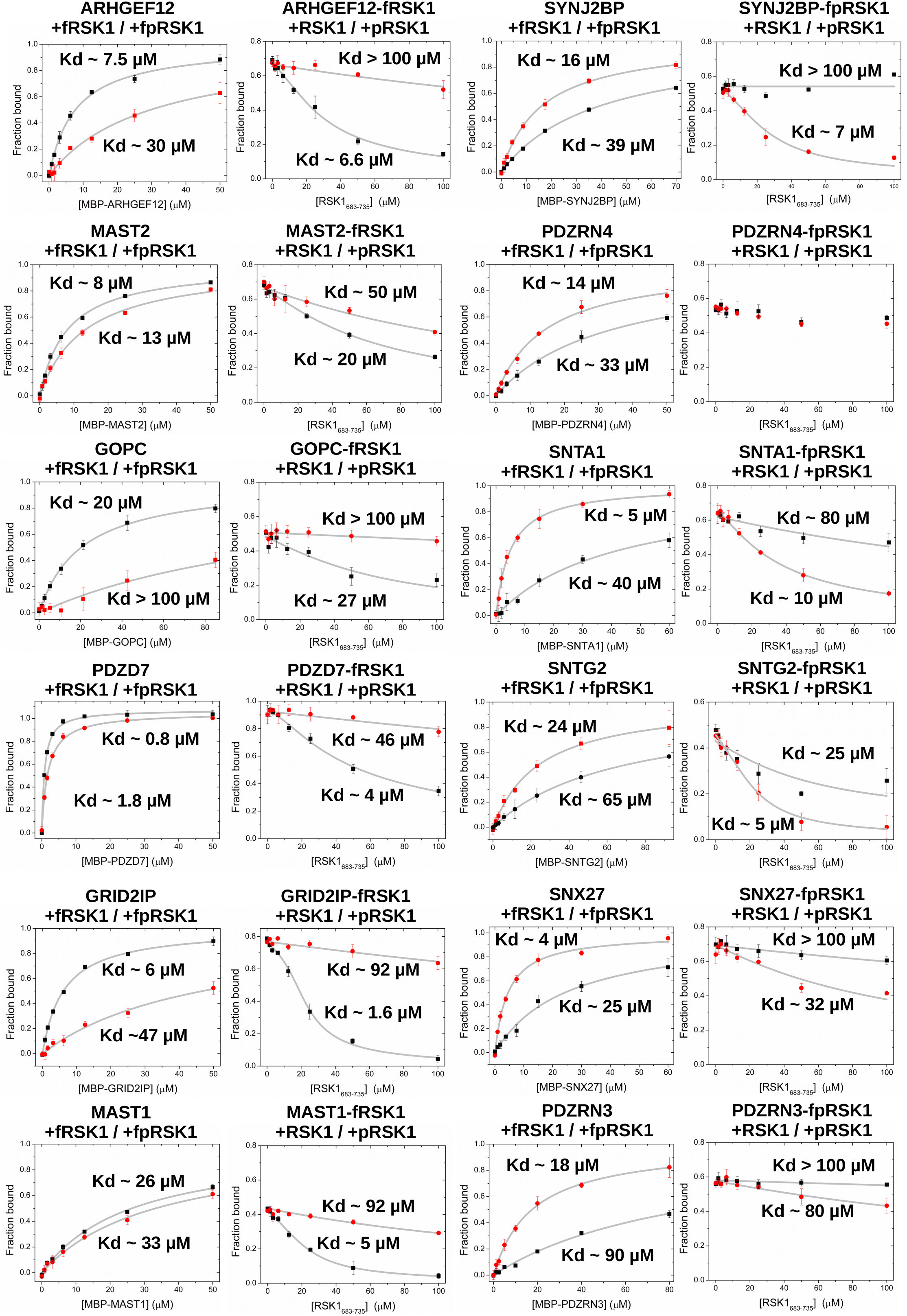
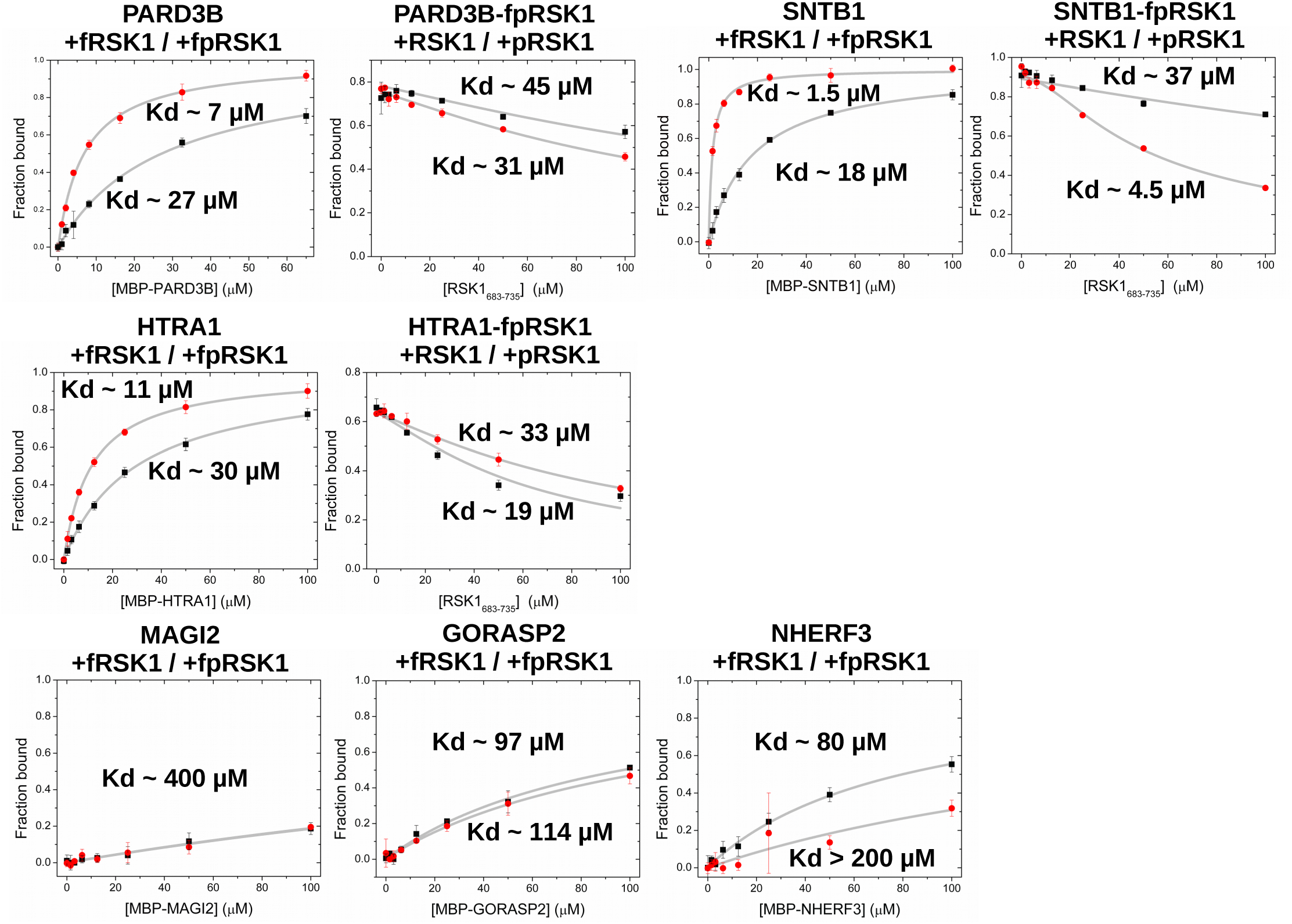
Results of the fluorescence polarization measurements. Fluorescence polarization measurements were carried out to measure the binding of multiple PDZ domains. Direct binding was measurement with a fluorescein labeled 7 residue long RSK1 peptide, while competitive measurements were measured with a 40 residue long native or monophosphorylated peptide (colored black and red, respectively).

**Figure S5.**
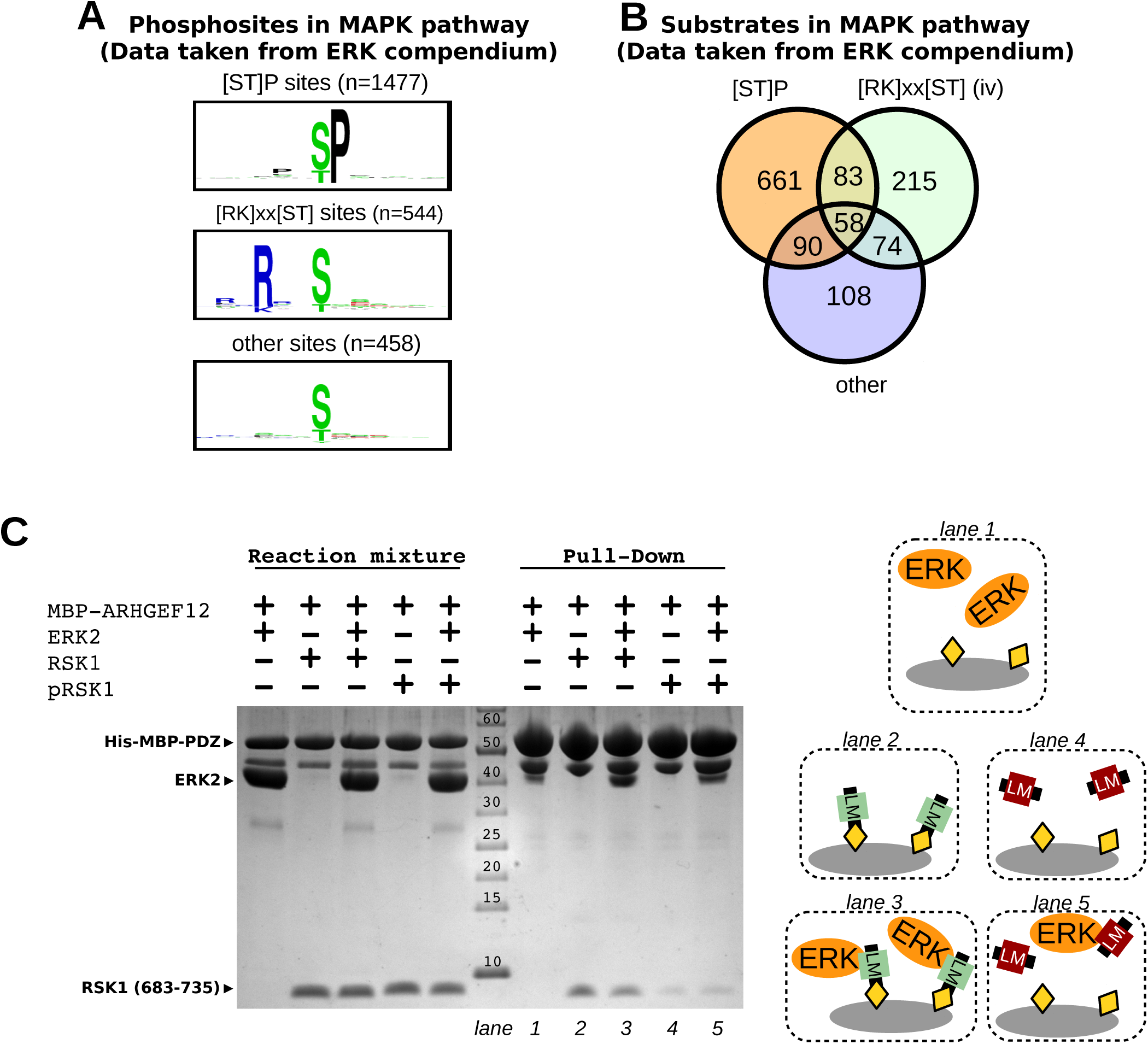
ERK cooperates with RSK. (A) We filtered the ERK compendium to [ST]P (phosphorylated directly by ERK), [RK]×[ST] (phosphorylated indirectly by RSK) and other (phosphorylated indirectly by an unknown kinase) phosphosites. We have found that only 60% of the phosphorylation sites were potential direct ERK substrates and 20% of the identified phosphorylation events were potential RSK substrates. (B) Most substrate proteins were phosphorylated on multiple sites by a single kinase, but there is a significant overlap between the different phosphorylation motifs indicating that a fraction of the substrates can be phosphorylated by more than one kinase. We considered these putative RSK substrates as an individual set of potential substrates in further analyses. (C) Ternary complex formation between RSK, ERK and a PDZ domain. The MBP-PDZ domain was used as a prey and RSK peptides, ERK2 or ERK2 bound RSK peptides were used as baits in a pull-down experiment. Proteins were detected on an SDS-PAGE gel stained with Coomassie protein dye. The panel shows the results of a representative MBP pull down assay from two independent experiments. We were able to detect an enhanced interaction between ARHGEF12 and ERK2 in the presence of unphosphorylated RSK1 peptide, confirming a ternary complex formation. Moreover, the phosphorylated RSK peptide was unable to induce such effect, confirming the OFF dimmer effect of the complex formation in this particular interaction.

**Figure S6.**
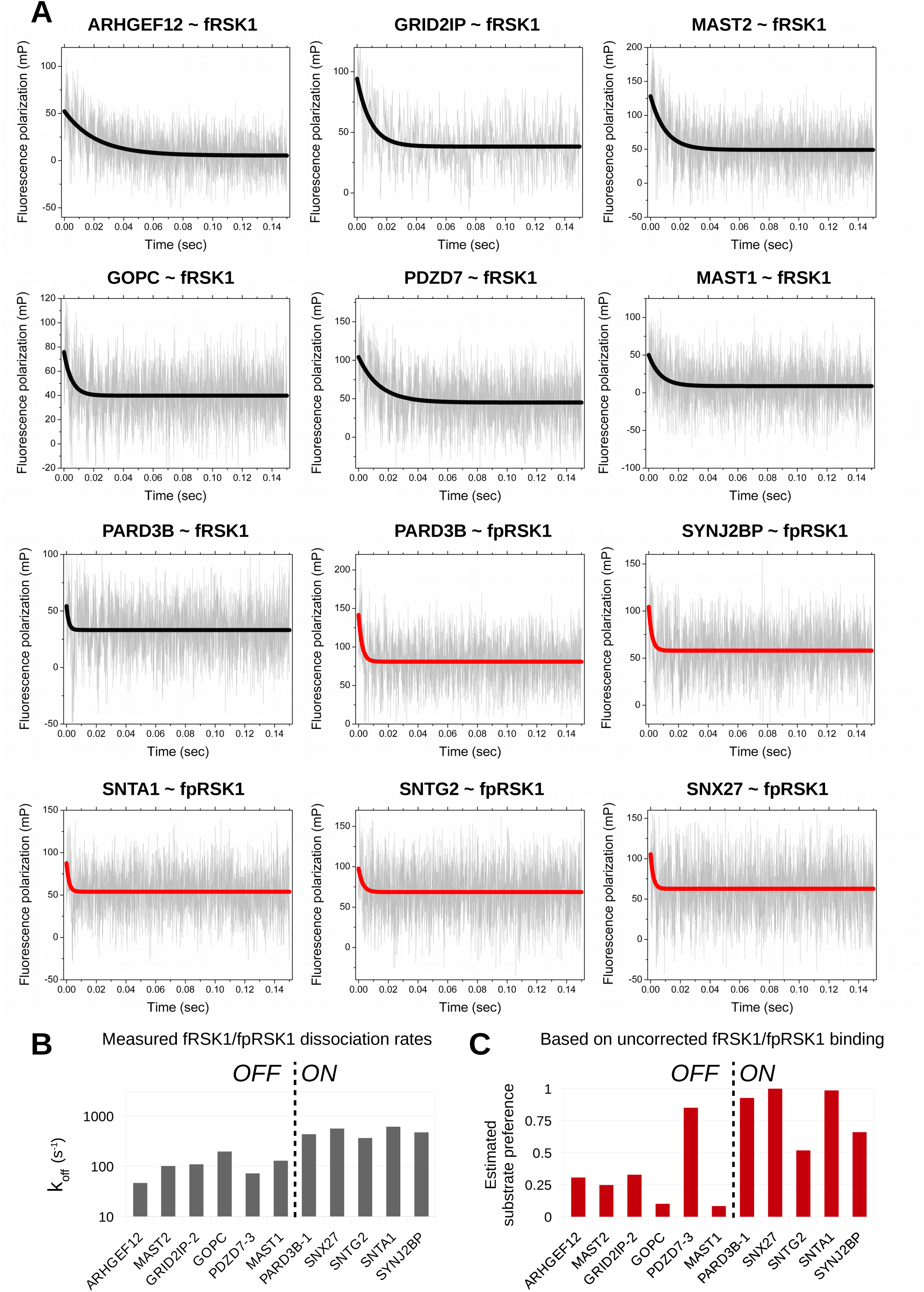
Summary of the stopped flow measurements. (A) A complexed fluorescent RSK1 peptide was mixed with high amount of unlabeled peptide. The change in the fluorescence polarization was monitored during the dissociation phase. (B) Measured off-rates of the labeled peptides. (C) Substrate phosphorylation was *in silico* estimated using their measured dissociation rate.

**Figure S7.**
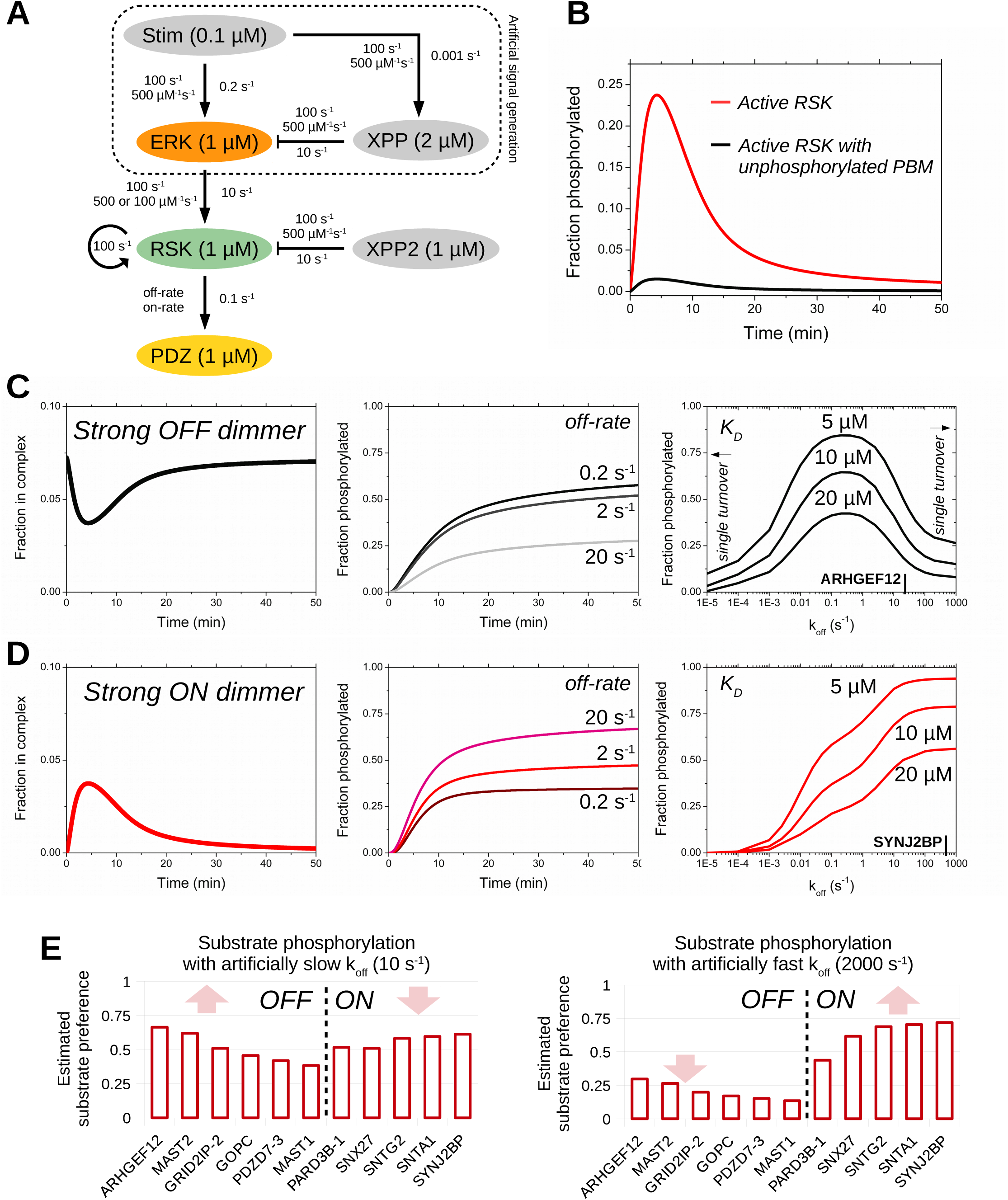
*In silico* modeling of PDZ substrate phosphorylation by RSK1. (A) This simplified mathematical model was used to simulate MAPK pathway activation. (B) Network based simulation shows that only a small fraction of activated RSK1 has an unphosphorylated PBM (even in the presence of high amount of PDZ domain). (C) Interaction partners with negative feedbacks show a dissociation upon stimulation. While the dissociation profile is off-rate dependent, the substrate phosphorylation rate is not. The system shows an optimal substrate phosphorylation at a low dissociation rate. (D) In contrast to the OFF dimmers, substrates with a positive feedback show an association profile. Increasing their dissociation kinetics increases their substrate phosphorylation rate. Note that the dynamical profiles of the interactions are very similar to the results of our cell based measurements, but we do not have any periodicity in this isotropic system. (E) A set of RSK substrates were *in silico* phosphorylated using an artificially slow or fast dissociation rate. Partners, which showed an OFF dimmer behavior preferred a slower binding kinetics while the ON dimmers preferred faster kinetics.

**Figure S8.**
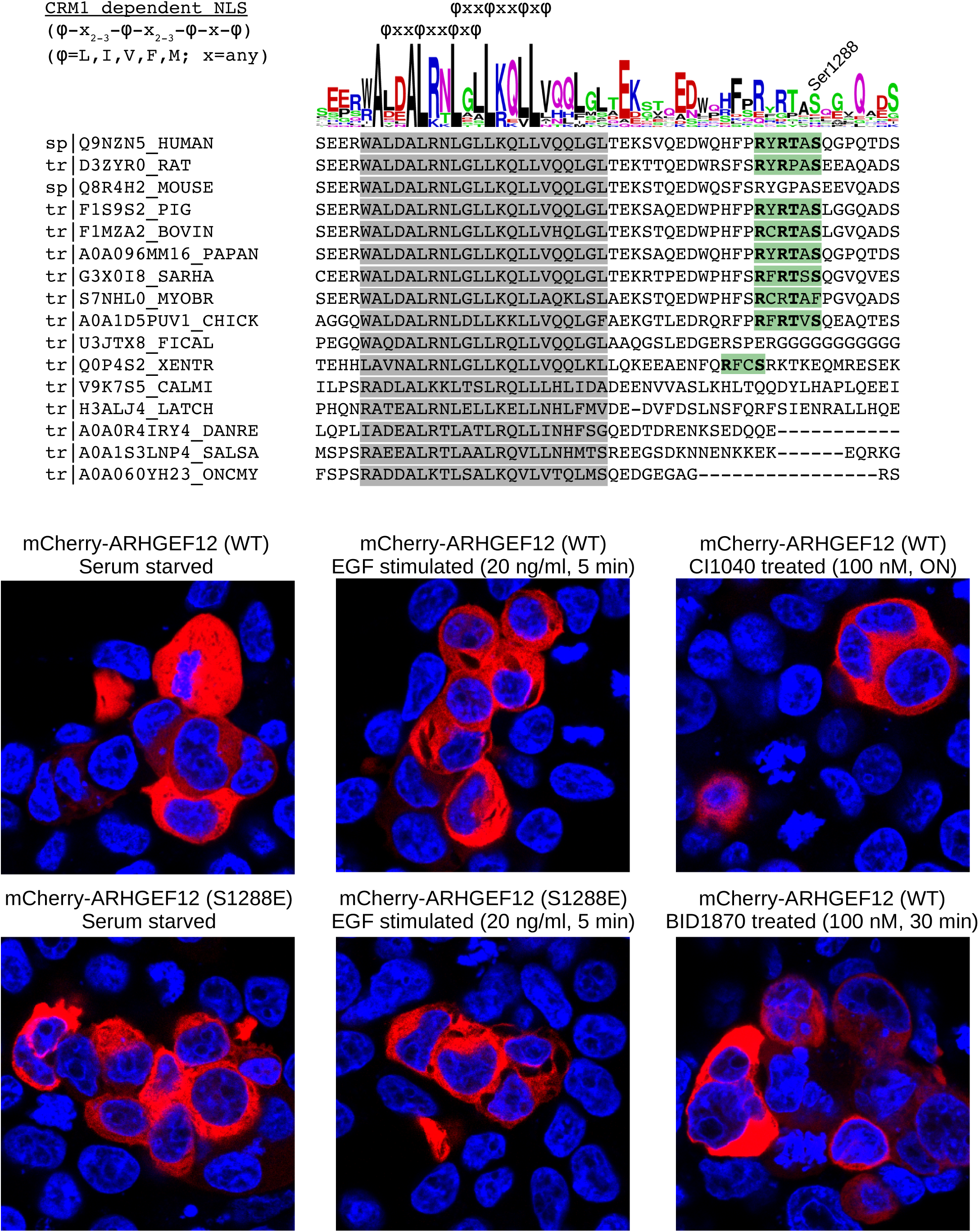
The unknown motif in ARHGEF12, next to the RSK phosphorylation site is probably not a phosphorylation induced NES. This conserved motif, which was previously described as a CRM1 dependent NES, can be found in most organisms and the RSK phosphosite lies next to it. Neither stimulation, phosphomimication or RSK/MEK inhibition affected the intracellular localization of the mCherry fused ARHGEF12 in HEK293T cells.

**Table S1.** Results of the HU assay. (BI values for both peptide.)

**Table S2.** The RSK substrate compendium

**Table S3.** PDZ scaffold mediated complexes

## Materials and methods

### Holdup assay

The automated holdup assay was carried out against peptides (RSK1_725–735_) in triplicates as previously described (Vincentelli *et al*, 2015) with minor modifications. For the detailed protocol please look at (Duhoo *et al*). The sequences of the clones of the PDZome v2 were designed according to (Luck *et al*, 2012). All genes were codon optimized for *E. coli* expression and cloned in a pETG41A plasmid. All protein constructs were expressed in *Escherichia coli* following the previous protocol (Vincentelli *et al*, 2015) with minimum modifications. All constructs were checked for solubility and cell lysate soluble fractions were adjusted to approximately 4 µM concentration and frozen in 96 well plates. Additionally, mass spectrometry was used to confirm the identity of each PDZ clones. For the detailed protocols of production and quality control, please look at (Duhoo *et al*). We measured interactions against 255 proteins with the unphosphorylated peptide and against 252 proteins with the phosphorylated peptide. In this work, BI = 0.2 was used as the minimal BI threshold value to define high-confidence PDZ-PBM pairs, as proposed previously (Vincentelli et al, 2015). Figure S1 contains the BI values of the RSK1 and phospho-RSK1 datasets. Data were analyzed as formerly described (Vincentelli *et al*, 2015). All plots and calculation in this work were calculated using these conventional datasets. Additionally, we already provide the values calculated with an updated protocol in the supplemental file, because the new calculation approach will set the standard for future holdup papers. These were generated using an automated informatic protocol awaiting for publication. This updated analysis revealed three new interaction partners of the native RSK1 peptide (SCRIB-3, MPDZ-10 and RHPN1) and four new partners of the phosphorylated peptide (SCRIB-3, LIN7A, PDZRN3-2 and DLG3). Apart from these weak interaction partners, most values are extremely coherent between calculations.

### Protein expression and purification

Tandem affinity (Ni-and MBP-) purified MBP-PDZ proteins were used in biochemical assays. ERK2 was expressed with an N-terminal cleavable hexahistidine tag which was removed after affinity purification. The affinity purified kinase was ion exchanged on a Resource Q, anion exchange column.

### Peptide synthesis

Unphosphorylated RSK1_683–735_ peptides were recombinantly expressed with an N-terminal cleavable GST tag. After affinity purification, the GST tag was removed and the peptide was isolated by reverse phase HPLC. A fraction of the isolated peptide was phosphorylated with a constitutively active (T573E mutant) RSK1 C-terminal kinase domain as formerly described (Vincentelli *et al*, 2015). Unphosphorylated, phosphorylated, fluorescein labeled or unlableled RSK1_729–735_ peptides were all chemically synthesized on an automated PSE Peptide Synthesizer (Protein Technologies, Tucson, AZ, USA) with Fmoc strategy. Biotinylated RSK1_725-735_ peptides were purchased from JPT Innovative Peptide Solutions with 70-80% purity. The biotin group was attached to the N-terminal via a TTDS linker. Protein (and Tyr containing peptide) concentrations were determined by UV spectroscopy. For peptides that lacked an aromatic residue, we directly measured their dry mass. Predicted peptide masses were confirmed by mass spectrometry.

### Isothermal titration calorimetry (ITC)

ITC measurements were carried out in 20 mM Hepes pH 7.5, 150 mM NaCl, 500 µM TCEP using a VP-ITC apparatus (MicroCal). 50 µM MBP-PDZ domain was titrated with concentrated peptides at 37 °C. The Origin for ITC 5.0 (Originlab) software package was used for data processing.

### Surface plasmon resonance (SPR)

SPR measurements were performed on a Biacore T200 instrument equipped with CM5 sensor chip. Streptavidine was immobilized on the sensor chip with EDC-MS using a standard protocol. Biotinylated peptides (RSK1, pRSK1, HPV16E6) were immobilized on streptavidine and after an extensive washing step, MBP-PDZ domains were injected onto the chip at 8 different concentrations and with two additional replicates. Unfortunately, our SPR analysis did not reveal the kinetic parameters of the studied PDZ-peptide interactions due to biphasic and very fast behavior. The saturated phase of the reference channel subtracted data was fitted with a hyperbolic function.

### Steady state fluorescence polarization

Fluorescence polarization was measured in 384-well plates (Corning) using Synergy H4 multi-mode reader (BioTek). For direct titration experiments, 50 nM reporter peptide (RSK1_729-735_) was mixed with increasing amount of MBP-PDZ domains. In competitive measurements, the 50 nM reporter peptide was mixed with the PDZ domain in a concentration to achieve high degree of complex formation. Subsequently, increasing amount of unlabeled peptide (RSK1_683-735_) was added to the reaction mixture. Titration experiments were carried out in triplicate and the average FP signal was used for fitting the data to a quadratic or competitive binding equation.

### Monte Carlo modeling

To estimate the dissociation constant of weak interactions, we used the measured BI values from the HU assay. This parameter equals the bound fraction of the PDZ domain, therefore, it can be inserted directly into the general binding equation:

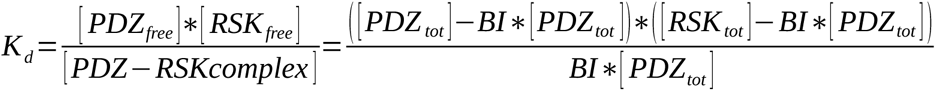
Assuming that the total PDZ domain concentration is ∼ 4 µM, the only unknown parameter is the total peptide concentration. Instead of a simple nonlinear fit, we have used Monte Carlo modeling to utilize the standard deviations of the HU assay and the K_d_ measurements. Each fitting was repeated 10000 times and the average peptide concentration along with the lower and upper quartiles were plotted in figure 3A. Based on our SPR measurements, the RSK peptide concentration should be around 20 µM (most probably between 18-21 µM). Direct FP indicates that this concentration should be around 14 µM (most probably between 6-20 µM). In the case of the competitive FP, we have found that the peptide concentration should be around 14 µM (most probably between 9-18 µM). For K_d_ extrapolation, we have used a peptide concentration of 17 µM.

### Protein-protein interaction assay

The NanoBiT PPI MCS starter system was purchased from Promega. Full-length RSK1 was cloned into pBit2.1-N[TK/SmBiT] vector. Full-length MAGI1 and ERK2 constructs were previously cloned into the LgBiT vector. Full-length ARHGEF12 (isoform 2), GOPC (isoform 2), PARD3B (1-913) and SYNJ2BP were cloned into the pBit1.1-N[TK/LgBiT] vectors. All constructs were cloned from HEK293T or HeLa cDNA pools and were confirmed by sequencing. HEK293T cells were cultured in Dulbecco’s Modified Eagle Medium (DMEM, Lonza) containing 10% fetal bovine serum and 1% penicillin/streptomycin/amphotericin B. 2×10^4^ cells/well were seeded onto a white, TC treated 96-well plate (Greiner) 24 hours prior to transfection. Transient transfections were carried out with FuGene HD reagent (Promega) according to the NanoBiT system’s instructions. 4 hours after transfection, cells were starved for 20 hours in CO_2_-independent medium (Thermo). Cells were assayed 24 hours after transfection using Nano-Glo reagent (Promega) and a Synergy H4 plate reader (BioTek). Experiments were carried out according to the manufacturer’s instructions. Stimulation was performed using 20 ng/ml EGF (Sigma-Aldrich). Each experiment was performed with at least 6 biological replicates. We must note that it is likely that the periodicity is very environment dependent and under slightly modified conditions (i.e. CO_2_ incubator instead of CO_2_ independent cell medium, different cell density or protein expression level…) no periodic features appeared.

### Pull-down experiment

For the *in vitro* pull-down experiment, Amylose resin (New England BioLabs) was saturated with Ni-NTA purified MBP-ARHGEF12. The immobilized PDZ domain was incubated with 60-60 µM full length ERK2 and RSK1 peptides in 500 µl (10-20 column volumes) for 30 min at room temperature. Amylose beads were separated by centrifugation and washed three times with 500 µl binding buffer (20 nM Hepes pH 7.5, 150 mM NaCl, 150 µM TCEP). Retained proteins were eluted with SDS loading buffer and the samples were subjected to TRIS-Tricine SDS-PAGE and stained with Coomassie protein dye.

### Signaling pathway modeling

Rule based network modeling was carried out with the software package BioNetGen with the ordinary differential equation solver running on a desktop PC. The simulated pathway was described in figure S7A. Each pathway activation was initiated from a pre-equilibrated state. The simulation was initiated by introducing the “Stim” to the system. This simplified, artificial signal generator was adjusted to mimic the natural activation profile of the ERK pathway upon EGF stimulation.

### Stopped-flow fluorescence polarization

Fast kinetic measurements were performed with the stopped-flow instrument SFM-300 (Bio-Logic) with polarized excitation at 488 nm. Parallel and perpendicular fluorescent emission were measured through a 550 +/-20 nm band pass filter (Comar Optics). All reactions were measured at 25 °C in a buffer containing 20 nM Hepes pH 7.5, 150 mM NaCl, 150 µM TCEP. Post-mixing fluorescent peptide concentration was 0.5 µM. The fluorescent peptide (RSK1_729-735_) was pre-complexed with high amount of MBP-PDZ domain (5-40 µM, post-mix). To measure the dissociation of the labeled peptide, we rapidly mixed the PDZ bound complex with high molar excess of unlabeled peptide (RSK1_729-735_ 100 µM, post-mix). Each experiment was carried out multiple times (n>9) and the averaged transients were fitted using a single exponential function. Corrections were applied to estimate the unbiased binding of an unlabeled peptide based on the dissociation constant differences between the direct FP measurements and the unbiased HU assay.

### Immunofluorescence

For detection of the intracellular localization of transfected proteins 1×10^5^ cells/well were seeded onto a cover slip-containing (Assistent) 24-well plate. Cells were fixed with 4% PFA solution and blocked for 1 hour in 5% BSA and 0.3% Triton-X 100, dissolved in PBS at room temperature. The RSK1/2 knockout (CRISPR) HEK293 cell line was a kind gift from Fanxiu Zhu. To introduce exogeneous WT or mutant RSK1 into these cell lines, we created pIRES2-EGFP based vectors, which expressed untagged RSK1s along with a GFP transfection reporter gene. Phosphorylated RSK was detected with the help of anti-pRSK pSer380 (1:800, CST) primary and Alexa Fluor 647 (anti-rabbit, 1:800, Thermo) conjugated secondary antibodies. ARHGEF12 (isoform 2) was cloned into a pmCherry-C1 vector. Mutations were introduced by quick change mutagenesis. Nuclear staining was performed using DAPI (0.1 μg/ml). After washing, cover glasses were mounted to microscopy slides by Mowiol 4-88 mounting medium (Sigma). Confocal microscopy was carried out using a Zeiss LSM 710 system (Carl Zeiss Microscopy GmbH, Jena, Germany) with a 40X oil objective. Images were processed by the ImageJ software.

### RhoA activation assay

The commercially available luminescence based G-LISA RhoA activation assay (Cytoskeleton) was used to measure the GTP bound RhoA levels in cell cultures. 2×10^5^ cells/well were seeded onto a 24-well plate. G-LISA assay was performed according to the manufacturer’s recommendations 24 hours after transfection with the exception of the concentration and the antibody dilutions. Sample concentrations were equalized to 1 mg/ml. Primary and secondary antibodies were diluted to 1:500 and 1:1000, respectively. Luminescence signal was detected on a Synergy H4 plate reader (BioTek). The RSK inhibitor BI-D1870 treatment was performed at 100 nM for 1h. The MEK inhibitor CI1040 was incubated ON at 100 nM. Inhibitor treatments were performed in Dulbecco’s Modified Eagle Medium supplemented with 10% fetal bovine serum. Serum stimulation (20%) was performed with serum starved cells for 5 min.

## Acknowledgements

This work was supported by the National Research Development and Innovation Office, Hungary (NKFIH): K 119359 (to LN), NN114309 and KKP 126963 (to AR). GG and MS were supported through the New National Excellence Program of the Hungarian Ministry of Human Capacities. The work was supported in part by grants from Ligue contre le Cancer (équipe labellisée 2015 to G.T.), National Institutes of Health (grant R01CA134737 to G.T.) and from European Union (PDZnet network, Marie Sklodowska-Curie grant No 675341 to G.T. and P.J.) This work was supported by the French Infrastructure for Integrated Structural Biology (FRISBI) ANR­10­INBS­ 05.

## Author contributions

GG conceived the project, carried out the experiments, analyzed data, and wrote the paper. LN, GT supervised the research, analyzed data, and wrote the paper. FB, PJ, YN, RV and GT performed and analyzed the holdup experiments. BB-K, CK, VB and MS contributed by carrying out cell-based and in vitro experiments.

